# Rapid turnover of CTLA4 is associated with a complex architecture of reversible ubiquitylation

**DOI:** 10.1101/2023.12.31.573735

**Authors:** Pei Yee Tey, Almut Dufner, Klaus-Peter Knobeloch, Jonathan N. Pruneda, Michael J. Clague, Sylvie Urbé

## Abstract

The immune checkpoint regulator CTLA4 is an unusually short-lived membrane protein. Here we show that its lysosomal degradation is dependent on ubiquitylation at Lysine residues 203 and 213. Inhibition of the v-ATPase partially restores CTLA4 levels following cycloheximide treatment, but also reveals a fraction that is secreted in exosomes. The endosomal deubiquitylase, USP8, interacts with CTLA4 and its loss enhances CTLA4 ubiquitylation in cancer cells, mouse CD4^+^ T cells and in cancer cell-derived exosomes. Depletion of the USP8 adapter protein, HD-PTP, but not ESCRT-0 recapitulates this cellular phenotype, but shows distinct properties *vis-à-vis* exosome incorporation. Re-expression of wild-type USP8, but neither a catalytically inactive, nor a localisation-compromised ΔMIT domain mutant can rescue delayed degradation of CTLA4, or counteract its accumulation in clustered endosomes. UbiCRest analysis of CTLA4-associated ubiquitin chain linkages identifies a complex mixture of conventional Lys63- and more unusual Lys27- and Lys29-linked polyubiquitin chains that may underly the rapidity of protein turnover.

## Introduction

The emergence of immunotherapy has dramatically changed the oncology landscape (Sanmamed and Chen, 2018). The cell surface receptor, CTLA4 (cytotoxic T-lymphocyte associated protein 4, CD152), provided the first such druggable immune checkpoint protein, offering positive outcomes in mouse models and clinical trials with CTLA4 blocking antibodies (Hodi et al., 2010; Kwon et al., 1997; Leach et al., 1996; Robert et al., 2011). Ipilimumab is a CTLA4 neutralising antibody approved as a monotherapy for advanced melanoma, whereas a second CTLA4 antagonist, Tremelimumab, is approved for combination use with other checkpoint inhibitors in cases of liver cancer (de Castria et al., 2022; Hargadon et al., 2018; Korman et al., 2022). Whilst showing robust and durable immune protection in cancer patients, the clinical benefits of CTLA4 blockade are often overshadowed by severe immunotherapy-related adverse events (irAEs) that highly resemble autoimmune reactions. Despite the clinical prominence of CTLA4, critical aspects of its cell biology are under-developed. In distinction to other immune checkpoint molecules (e.g. PD-L1 and PD-1), it has one of the shortest known half-lives amongst transmembrane proteins (Li et al., 2021; Rusilowicz-Jones et al., 2022). A full understanding of its turnover may lead to novel therapeutic strategies.

CTLA4 is expressed constitutively in a subset of immunosuppressive regulatory T cells (T_reg_), while being induced in activated CD8^+^ and CD4^+^ T cells to limit and terminate T cell signalling (Linsley et al., 1992; Takahashi et al., 2000). Upon T cell activation, an intracellular pool of CTLA4 is mobilised to the cell surface where it displaces T cell co-receptor CD28 for interaction with their shared ligands on antigen presenting cells (CD80 and CD86) (Linsley et al., 1996). It is now evident that CTLA4 expression is not exclusive to T cells; various tumours are reported to express CTLA4 and can also release CTLA4-containing exosomes (Contardi et al., 2005; Laurent et al., 2013; Mo et al., 2018; Paulsen et al., 2017; Theodoraki et al., 2019; Whiteside, 2013).

The dynamics of CTLA4 localisation are strongly linked to physiological function, yet the molecular mechanisms governing its fate are incompletely understood. It is rapidly internalised from the cell surface in a clathrin- and dynamin-dependent manner to create a majority intracellular pool at steady-state (Follows et al., 2001; Qureshi et al., 2012). T cell-associated CTLA4 has been shown to capture CD80 ligands from adjacent antigen-presenting cells *in vivo* for subsequent delivery to the lysosomes (Hou et al., 2015; Qureshi et al., 2011). Furthermore, CTLA4 is turned over by lysosomal degradation, displaying a short half-life of ∼2 hrs in activated mouse transgenic T cells and ∼3 hrs in CTLA4-over-expressing CHO cells (Egen and Allison, 2002; Khailaie et al., 2018). In T cells, LPS Responsive Beige-Like Anchor Protein (LRBA) deflects CTLA4 from this degradative route towards recycling, so that patients with LRBA deficiency suffer from early onset autoimmune disorders (Alroqi et al., 2018; Charbonnier et al., 2015; Lo et al., 2015). Otherwise, molecular details are fairly sparse. It is known to bind to the clathrin adaptors AP2 and AP1 for endocytosis and transport from the trans Golgi Network (TGN) to lysosomes, respectively (Leung et al., 1995; Qureshi et al., 2012; Schneider et al., 1999; Shiratori et al., 1997). Several Rab GTPases regulate CTLA4 sorting, with Rab5 and Rab7 mediating CTLA4 internalisation and downstream degradation, Rab11 controlling CTLA4 recycling and Rab8 regulating CTLA4 transport to the cell surface (Banton et al., 2014; Janman et al., 2021).

Ubiquitylation frequently provides a critical sorting signal by which receptors interact with the ESCRT (endosomal sorting complexes required for transport) machinery and commit to lysosomal degradation (Clague and Urbe, 2017b; Piper et al., 2014). The receptors are then incorporated into luminal vesicles of multi-vesicular bodies (MVBs) before fusion with lysosomes (Futter et al., 1996). Two endosomal deubiquitylating enzymes (DUBs), USP8 (UBPY) and AMSH (associated molecule with the SH3 domain of STAM; STAM binding protein, STAMBP), interact with the ESCRT machinery and can differentially influence the fate of endocytosed receptors (Clague et al., 2012; Clague and Urbe, 2006). It is known that CTLA4 can be ubiquitylated, but not whether this ubiquitylation influences endosomal sorting. Some circumstantial evidence supporting this hypothesis is provided by the induction of CTLA4 ubiquitylation upon engagement of its ligand, the transmembrane protein CD80, which is targeted for destruction by trans-endocytosis (Kennedy et al., 2022). Here, we first show that CTLA4 is endogenously expressed as a short-lived protein in selected cancer cell lines, where it is degraded in lysosomes. We demonstrate that direct ubiquitylation at two specific lysine residues controls CTLA4 lysosomal degradation. We uncover a critical role for USP8 and its adapter protein HD-PTP in regulating CTLA4 ubiquitylation status in multiple settings, whilst Ubiquitin Chain Restriction (UbiCRest) analysis reveals a distinctive architecture of CTLA4-associated polyubiquitin chains (Hospenthal et al., 2015).

## Results

### CTLA4 ubiquitylation promotes its rapid constitutive degradation

Individual reports indicate that a variety of tumour cells express endogenous CTLA4 (Chen et al., 2017; Laurent et al., 2013; Mo et al., 2018; Paulsen et al., 2017). We consulted the cancer cell line encyclopedia (CCLE) database for cancer cell lines expressing CTLA4, and selected one melanoma (A2058) and one squamous lung carcinoma (NCI-H520) cell line for further investigation (Figure **S1A**) (Barretina et al., 2012). Western blot analysis of total cell lysates prepared from these cells treated with non-targeting control (NT1) or CTLA4-selective siRNA, confirmed expression in both lines (Figure **S1B**). In parallel, we generated Flp-In HeLa cells that constitutively express an epitope-tagged form, CTLA4-HA. A cycloheximide time course established that CTLA4 has an extremely short half-life (< 1 h) in all three cell lines, which makes it one of the most short-lived transmembrane proteins described in human cancer cells (Figure **1A, B**) (Li et al., 2021; Rusilowicz-Jones et al., 2022).

**Figure 1.**
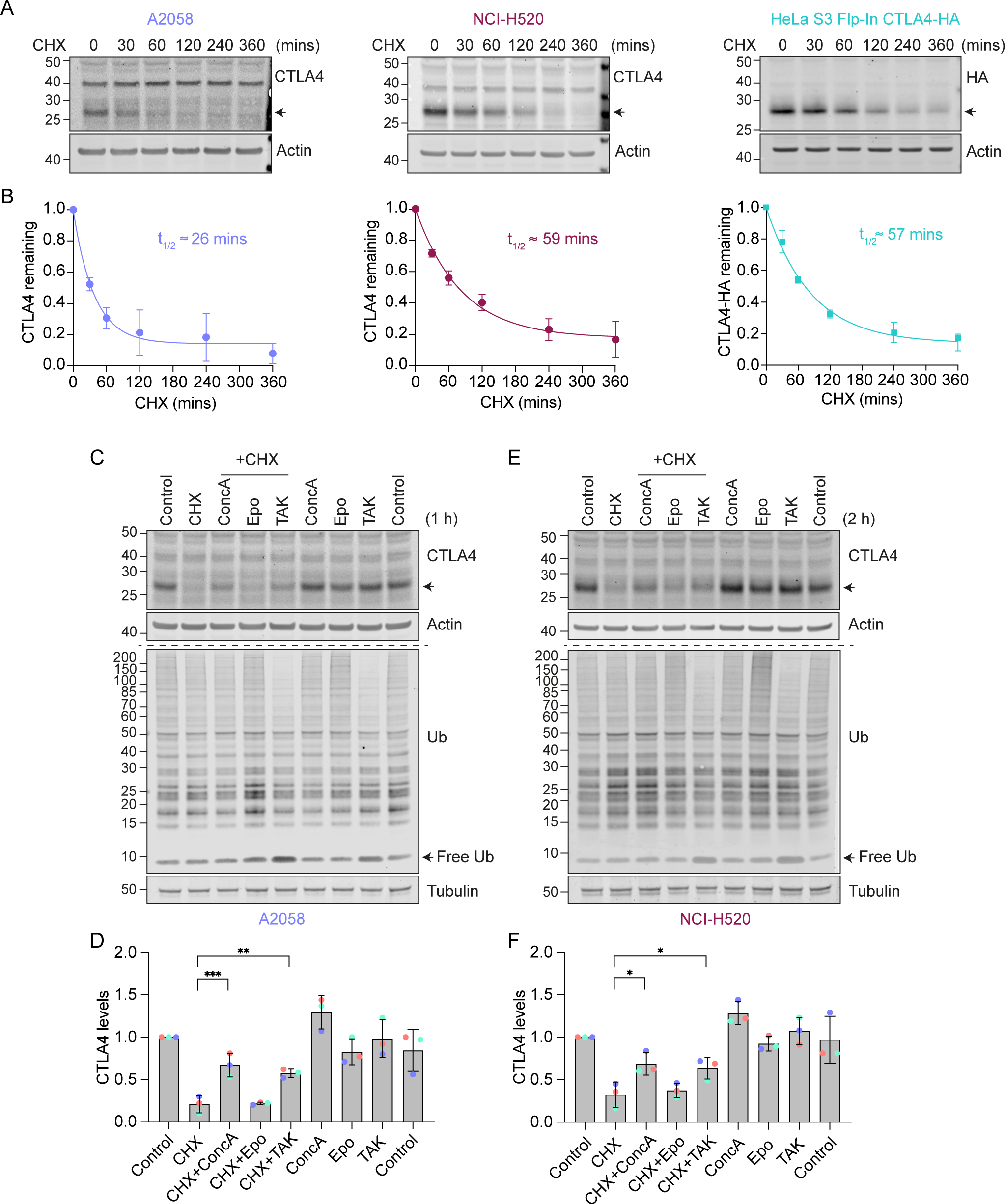
Lysosomal degradation of CTLA4 requires a functional ubiquitin conjugation cascade. **A.** Representative western blots of A2058, NCI-H520 and HeLa S3 Flp-In CTLA4-HA cells treated with Cycloheximide (CHX, 100 µg/ml) for indicated times prior to lysis. **B.** Quantification of data represented in (A). The half-life of CTLA4 in different cell lines was estimated using an exponential decay model. Error bars indicate standard deviation from three independent experiments. **C, E.** Representative western blots of A2058 and NCI-H520 cells treated with Concanamycin (ConcA, 100 nM), Epoxomicin (Epo, 100 nM) or TAK-243 (TAK, 1 µM) for 15 mins before the addition of CHX for 1 h (C, A2058) or 2 h (E, NCI-H520). **D, F.** Quantification of CTLA4 levels after treatment with indicated inhibitors relative to control for data represented in (C) and (E). Individual data points from three independent, colour-coded experiments are shown. Error bars show standard deviation. One-way ANOVA and Dunnett’s multiple comparison test , *p<0.05, **p<0.01, ***p<0.001.

We next set out to define the degradation route for CTLA4 in these cells using established inhibitors of the proteasome (Epoxomicin), lysosome (Concanamycin A; v-ATPase inhibitor) and ubiquitin conjugation machinery (TAK-243; UBA1 inhibitor). Both Concanamycin A (ConcA) and TAK-243 treatment rescued endogenous CTLA4 protein from degradation under Cycloheximide chase conditions in melanoma and lung cancer cells, whereas the proteasome inhibitor (Epo) was without effect (Figure **1C-F**). Probing the samples for ubiquitin reveals that proteasome and ubiquitin E1 inhibitors increase and decrease ubiquitin conjugates as expected. These data indicate that CTLA4 is constitutively targeted for lysosomal degradation in a ubiquitin-dependent fashion. This sorting pathway has been well-described for other transmembrane proteins, e.g. activated epidermal growth factor receptor (EGFR), and involves ubiquitin- and ESCRT machinery-dependent packaging of cargo into internal vesicles of MVBs (Henne et al., 2011; Hurley and Stenmark, 2011).

We noticed that in melanoma and lung cancer cells, blocking the lysosomal degradation pathway with ConcA only partially rescued constitutive CTLA4 degradation (Figures **1C-F** and **S2A**). We wondered whether a fraction of CTLA4 might not be degraded, but rather secreted via the release of exosomes (van Niel et al., 2018). These small extracellular vesicles are derived from the internal vesicles of specialised MVBs that can fuse with the plasma membrane and their release has previously been shown to be dramatically enhanced by v-ATPase inhibitors (Edgar et al., 2016). Harvesting conditioned media from A2058 cells treated for either 2 h or overnight (O/N) with ConcA, revealed secretion of established exosome markers CD63 and Syntenin (SDCBP) as well as CTLA4 (>20% of total, after O/N ConcA treatment) into the medium (Figure **2A, B** and **S2A**). In HeLa cells this pathway is less prominent; ConcA fully rescued CTLA4 protein levels in lysates of CHX treated cells, and an overnight treatment was necessary to visualise exosome and CTLA4-HA secretion (Figure **S2B-D**). A protease protection assay confirmed the expected orientation of CTLA4 in the exosome fraction, with an exposed amino-terminus (Figure **2C, D**).

**Figure 2.**
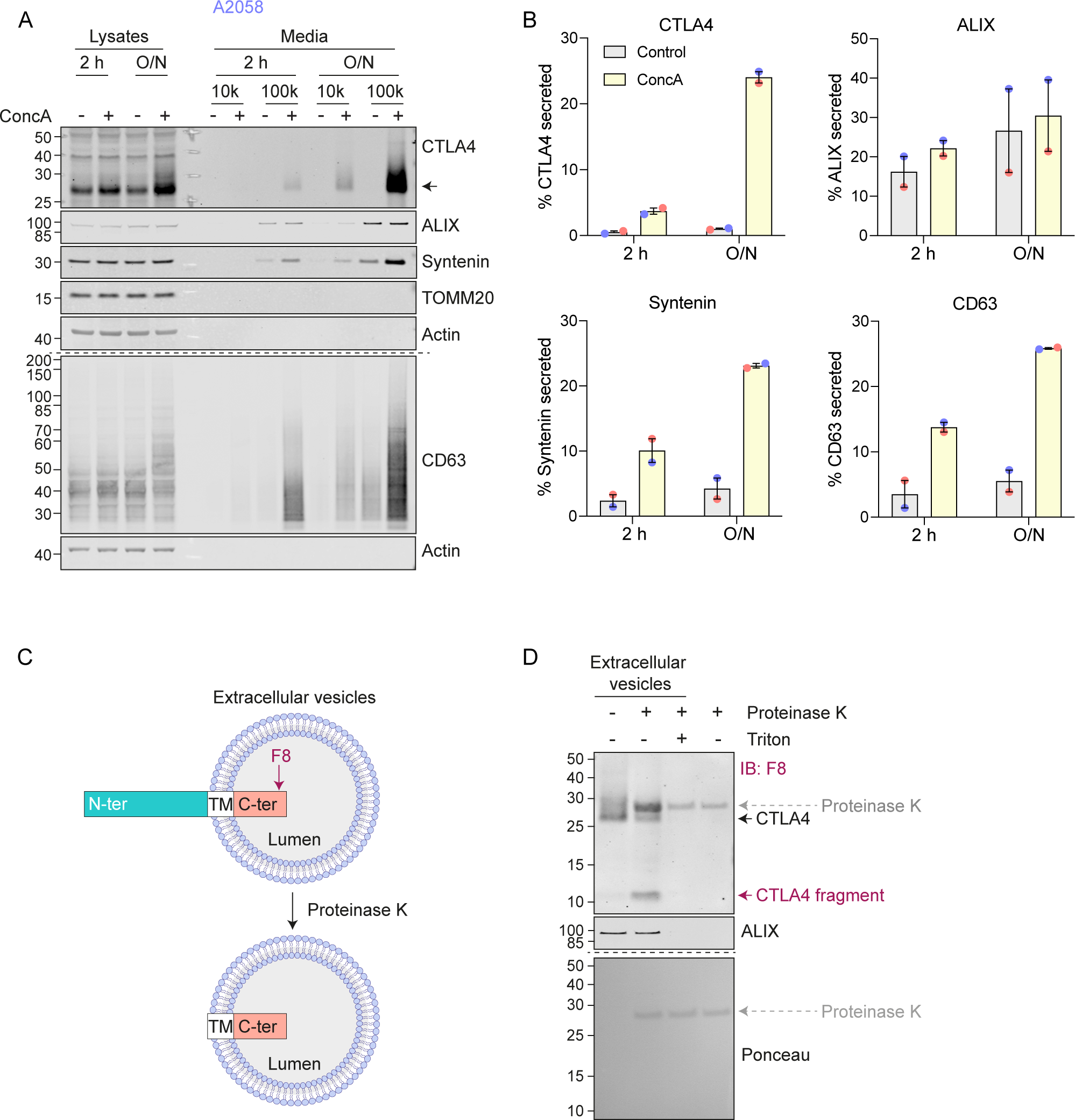
CTLA4 is secreted via exosomes in response to v-ATPase inhibition. **A.** Representative western blots of lysates and culture supernatants from mock- and Concanamycin A (ConcA, 100 nM) treated A2058 cells (O/N: overnight). Cells were lysed and the media collected by serial centrifugation to concentrate extracellular vesicles (100k pellet, exosome fraction). **B.** Quantification of CTLA4, Syntenin and CD63 recovered in the exosome fractions (100k) for data represented in (A). Individual data points from two independent, colour-coded experiments are shown. Error bars indicate range. **C.** Predicted topology of CTLA4 in exosomal membranes. F8: C-terminal (C-ter) cytoplasmic domain directed CTLA4 antibody; TM, transmembrane domain; N-ter, N-terminal domain. **D.** Representative western blots and Ponceau staining of samples from a Proteinase K protection assay of exosome-associated CTLA4. IB: Immunoblot.

### USP8 depletion enhances CTLA4 ubiquitylation but delays its degradation

Having established that ubiquitylation is required for lysosomal targeting and degradation of CTLA4, we reasoned that its trafficking may be regulated by endosome-associated DUBs, AMSH or USP8 (Clague and Urbe, 2006; Clague and Urbe, 2017). AMSH is a highly selective enzyme specialising in removing Lys63-linked ubiquitin chains, whereas USP8 cleaves a wide range of ubiquitin chain types (McCullough et al., 2006; Ritorto et al., 2014; Row et al., 2006). We used RNAi to deplete each of these DUBs and monitored the turnover of CTLA4 using a cycloheximide chase. Both endogenous and heterologuously expressed CTLA4(-HA) were insensitive to AMSH depletion (Figure **3A, B**). In contrast, siRNA targeting of USP8 increased CTLA4 half-life in both cell lines (Figure **3C, D**). We have previously shown that USP8 not only deubiquitylates endolysosomal cargo, but also stabilises the ESCRT-0 components HRS and STAM (Clague and Urbe, 2006; Row et al., 2006). Loss of HRS, upon USP8 depletion, is clearly apparent in both cell lines studied here (Figure **3C, D**). However, depletion of HRS itself had very little impact on CTLA4 turnover, thus establishing an ESCRT-0 independent role for USP8 in regulating CTLA4 degradation (Figure **3E, F**).

**Figure 3.**
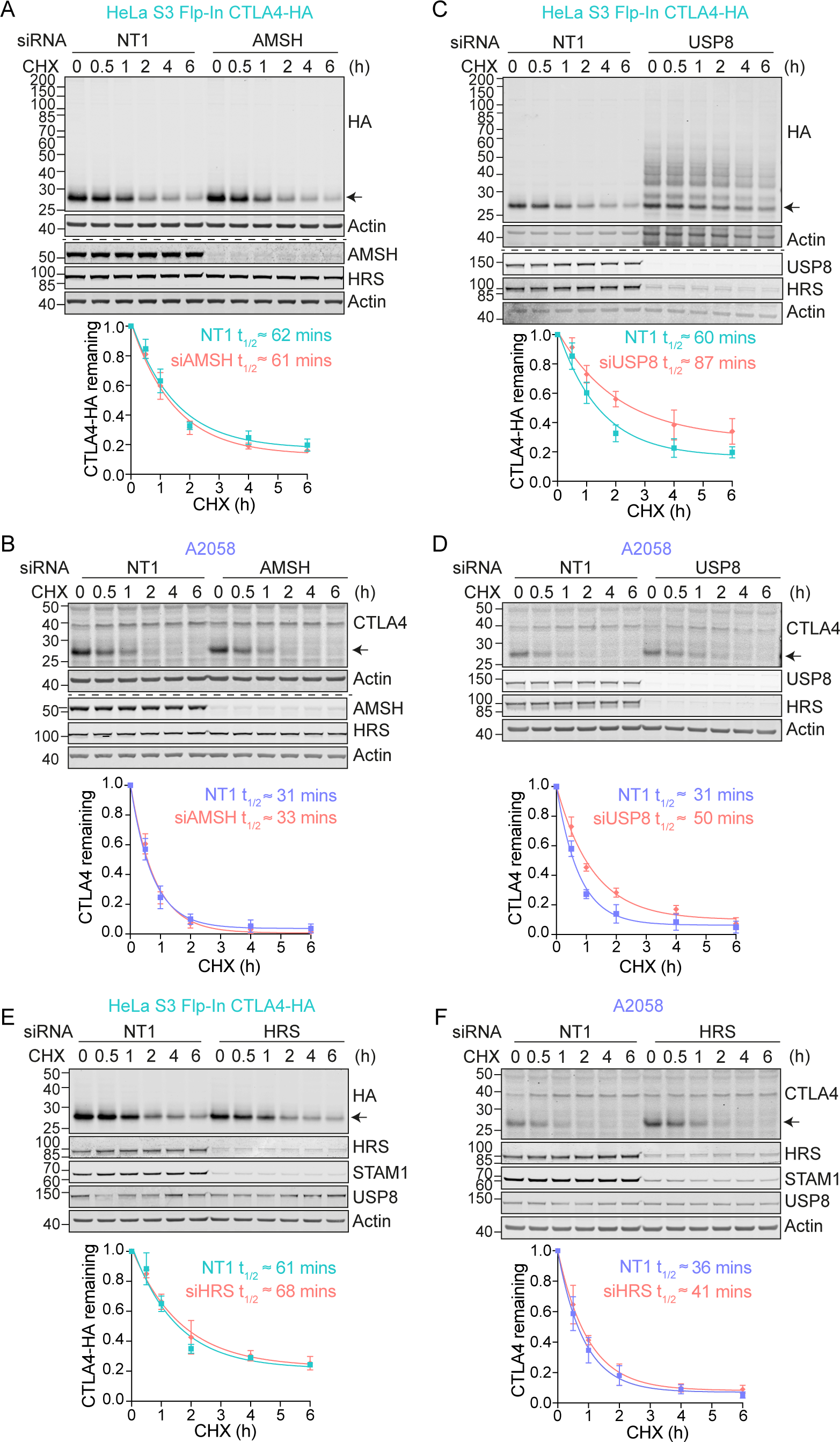
CTLA4 degradation is delayed in the absence of USP8. **A-F.** Representative western blots and associated quantifications of Cycloheximide (CHX; 100 µg/ml) chase experiments in HeLa S3 Flp-In CTLA4-HA (A, C) and A2058 (B, D) following treatment with non-targeting (NT1), AMSH, USP8 or HRS siRNA for 72 h. The half-life of CTLA4 was estimated using an exponential decay model. Error bars show standard deviation from three independent experiments.

In the absence of USP8, we noticed a series of higher molecular weight bands, indicative of ubiquitylated species of CTLA4 that were most apparent for CTLA4-HA (Figure **3C**). We next used two separate approaches to assess whether CTLA4 was ubiquitylated in these cells. First, we enriched ubiquitylated proteins using a Tandem Ubiquitin Binding Entities (TUBES) pulldown and probed for CTLA4 (Figure **4A, B**) (Mattern et al., 2019). Secondly we immunoprecipitated CTLA4-HA from denatured HeLa cell lysates and probed for ubiquitin (Figure **4C, D**). Together these experiments demonstrate that a fraction of CTLA4 is ubiquitylated at steady state, whilst depletion of USP8 but not AMSH dramatically increases this ubiquitylated pool. We also took advantage of a mouse model for conditional USP8 deletion (ΔUSP8). T cells derived from these mice can be cultured *in vitro* and treated with Tamoxifen to elicit USP8 deletion. It is immediately apparent that CTLA4 runs as a higher molecular species upon USP8 deletion in T cell populations isolated from three distinct sets of mice (Figure **4E**). A TUBES pulldown confirmed that these species correspond to ubiquitylated CTLA4.

**Figure 4.**
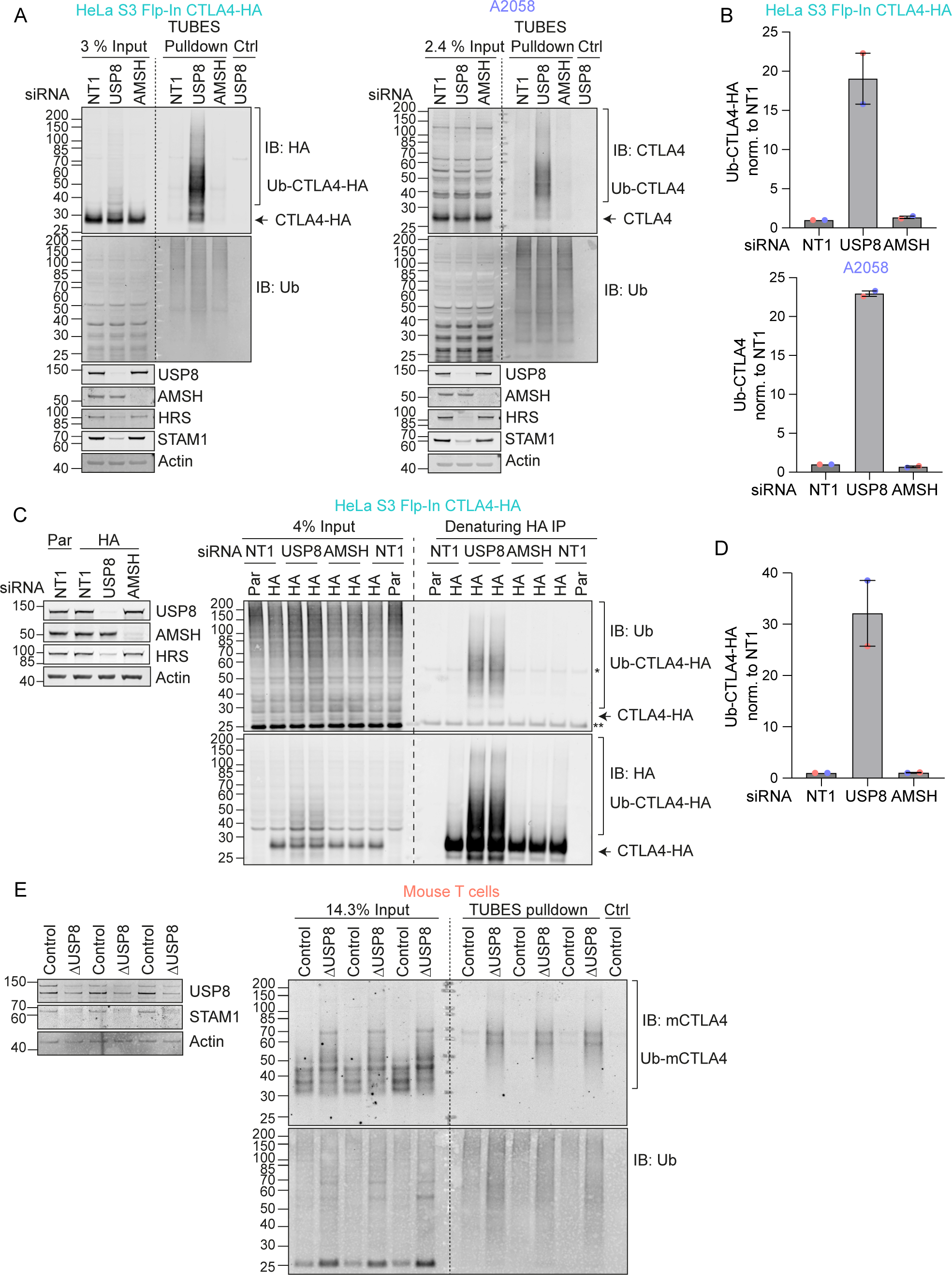
USP8 depletion dramatically enhances CTLA4 ubiquitylation. **A.** Representative western blots of TUBES pulldown of ubiquitylated CTLA4. HeLa S3 Flp-In CTLA4-HA and A2058 cells were transfected for 72 h with non-targeting (NT1), USP8 and AMSH siRNA prior to lysis. Lysates were subjected to TUBES-pulldown (IB: Immunoblot). **B.** Quantification of ubiquitylated CTLA4-HA isolated by TUBES pulldown for data represented in (A). Ubiquitylated CTLA4-HA was normalised to total ubiquitin pulled down. Individual data points from two independent, colour-coded experiments are shown. Error bars indicate the range. **C.** Representative western blots of ubiquitylated CTLA4-HA immunoprecipitated under denaturing conditions. HeLa S3 Flp-In parental (Par) or CTLA4-HA (HA) cells were transfected for 72 h with non-targeting (NT1), USP8 and AMSH siRNA. Cells were lysed in denaturing SDS lysis buffer and lysates subjected to immunoprecipitation (IP) with anti-HA coupled magnetic beads. *Antibody heavy chain; ** antibody light chain. **D.** Quantification of CTLA4-HA ubiquitylation relative to NT1 for data represented in (C). Ubiquitylated CTLA4-HA was normalised to total immunoprecipitated CTLA4-HA. Individual data points from two independent, colour-coded experiments are shown. Error bars indicate the range. **E.** Representative western blots of TUBES pulldown of ubiquitylated CTLA4. Lysates from USP8 fl/fl (Control, n=3) and USP8 deleted (ΔUSP8, n=3) activated T cells derived from individual mice were either analysed directly by SDS-PAGE and western blot (left blot), or first subjected to a TUBES pulldown prior to analysis alongside input samples.

### HDPTP and USP8 cooperate to govern CTLA4 ubiquitylation

Immunofluorescence microscopy of USP8-depleted HeLa and A2058 cells reveals the typical enlarged or clustered endosomal morphology as previously reported (Niendorf et al., 2007; Row et al., 2006). CTLA4 accumulates, on both EEA1 and LAMP1-positive structures marking early and late endosomes respectively (Figure **5A-H**). USP8 associates with multiple interaction partners at endosomal membranes, including STAM1 and 2, several CHMPs, HD-PTP (PTPN23) and ubiquitylated cargo itself (Clague and Urbe, 2006). Of these, HD-PTP has previously been shown to be essential for USP8 recruitment to activated receptors at endosomes (Ali et al., 2013; Kharitidi et al., 2015; Ma et al., 2015; Parkinson et al., 2021). We could detect a small fraction of endogenous USP8 associated with CTLA4-HA that was abolished by HD-PTP depletion (Figure **6A**). Consequently, loss of HD-PTP mirrors that of USP8, by promoting CTLA4-HA ubiquitylation and increasing its half-life (Figure **6A-G**). This further corroborates a role for HD-PTP as an obligatory adapter for USP8 recruitment. In A2058 cells the same effects of HD-PTP depletion upon ubiquitylation and half-life of endogenous CTLA4 are evident, albeit less strong than with USP8 depletion (Figure **6D-G**). As A2508 cells allow ready assessment of exosomal release we tested the effects of USP8 and HD-PTP depletion on this process. The ConcA induced release of CTLA4 is dependent on the exosomal sorting factors Syntenin and ALIX, whilst USP8 depletion enhances both the amount and ubiquitylation status of CTLA4 released into the media without altering the exosomal pool of Syntenin (Figure **S3A, B**). HD-PTP depletion is without effect on CTLA4-secretion, providing the first dissociation of USP8 and HD-PTP cellular phenotypes, perhaps reflecting another USP8 recruitment mode associated with this exosome directed aspect of USP8 function.

**Figure 5.**
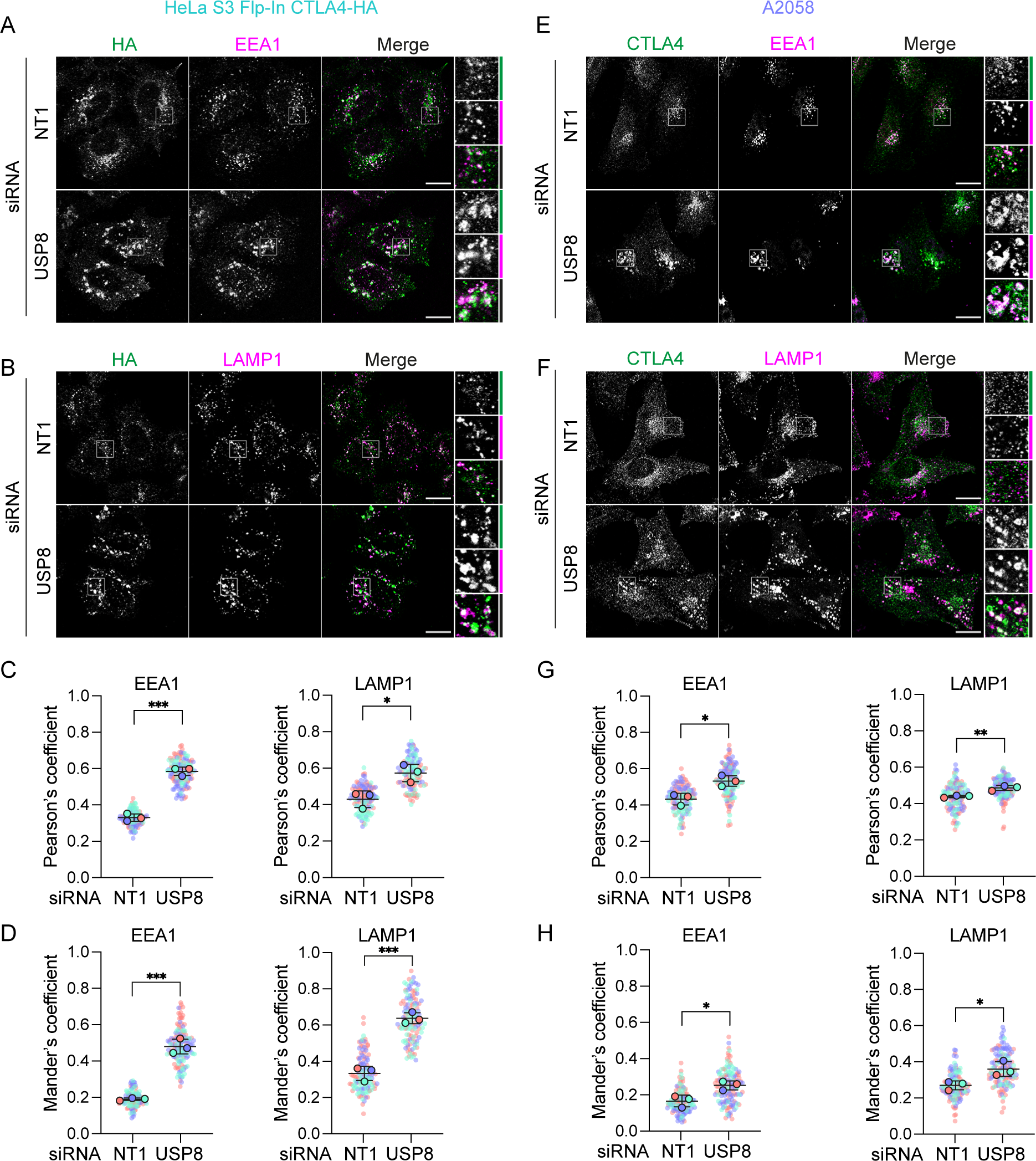
CTLA4 accumulates in enlarged endosomes following USP8 depletion. **A,B.** Representative confocal images of CTLA4-HA co-stained for EEA1 (A) or LAMP1 (B) in HeLa S3 Flp-In CTLA4-HA cells transfected for 72 h with non-targeting (NT1) or USP8 siRNA. Scale bar: 15 µm. **C,D.** Co-localisation analysis of CTLA4-HA with EEA1 and LAMP1 in HeLa S3 Flp-In CTLA4-HA for data represented in (A and B). Shown are the Pearson’s (C) or Mander’s coefficients (D). Error bars show standard deviation from three independent, colour-coded experiments. Opaque circles with black outlines correspond to the mean value from each experiment. Unpaired t-test. *p<0.05,***p<0.001 **E-F.** Representative confocal images of CTLA4 co-stained for EEA1 (E) or LAMP1 (F) in A2058 cells transfected for 72 h with NT1 or USP8 siRNA. Scale bar: 15 µm. **G-H.** Co-localisation analysis of CTLA4 with EEA1 and LAMP1 in A2058 cells for data represented in (E and F). Graphs show Pearson’s (G) or Mander’s coefficients (H) between CTLA4 and EEA1 or LAMP1. Error bars show standard deviation from three independent, colour-coded experiments. Opaque circles with black outlines correspond to the mean value from each experiment. Unpaired t-test. *p<0.05,**p<0.01

**Figure 6.**
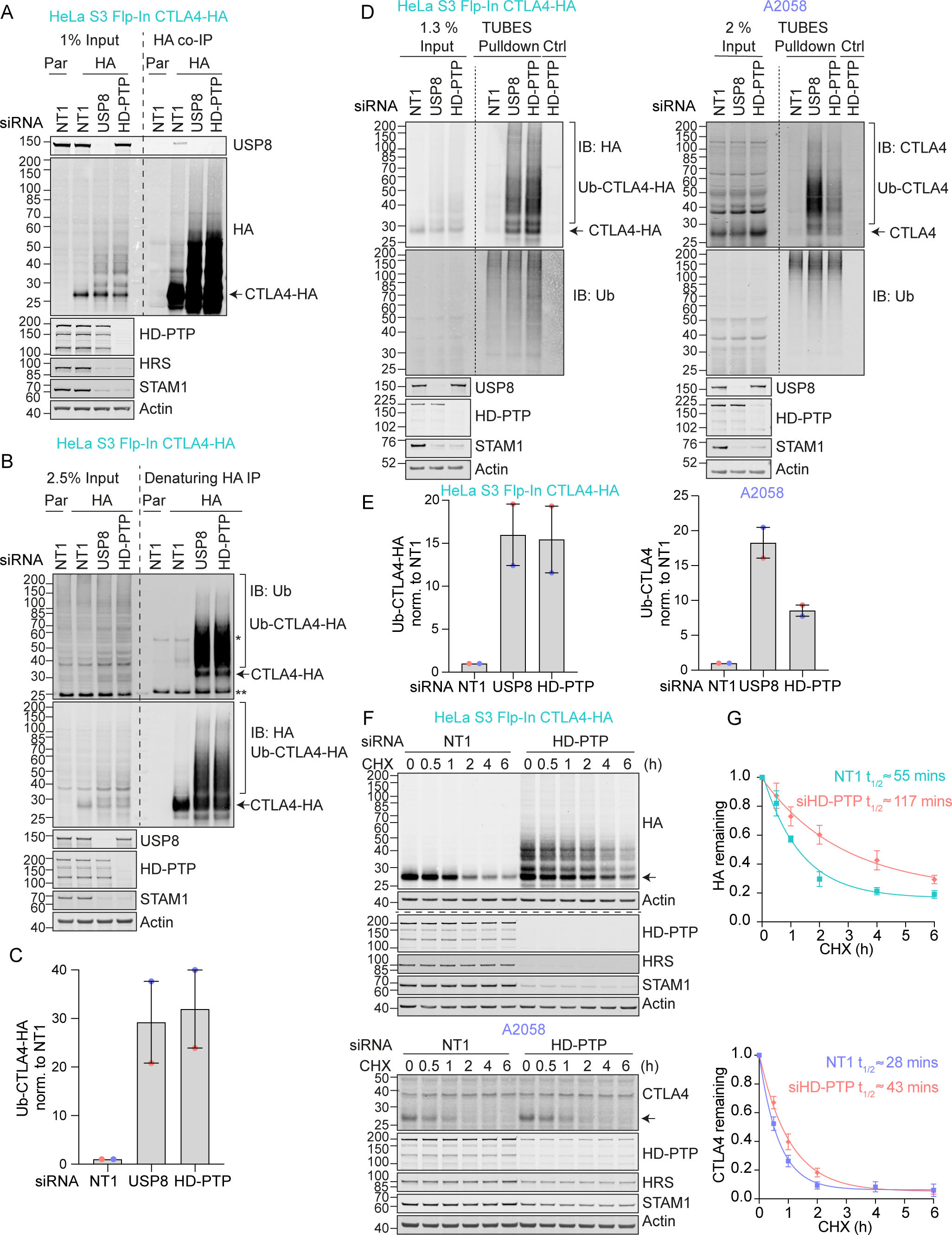
Regulation of CTLA4 ubiquitylation by USP8 is dependent on HD-PTP. **A.** Representative western blots of USP8 co-immunoprecipitated with CTLA4-HA. HeLa S3 Flp-In parental (Par) or CTLA4-HA (HA) cells were transfected for 72 h with non-targeting (NT1), USP8 or HD-PTP siRNA. Cells were lysed and lysates subjected to HA-immunoprecipitation (IP; IB: Immunoblot). **B.** Representative western blots of ubiquitylated CTLA4-HA immunoprecipitated under denaturing conditions. HeLa S3 Flp-In parental (Par) or CTLA4-HA (HA) cells were transfected as in (A). Cells were lysed in denaturing SDS lysis buffer and lysates subjected to immunoprecipitation (IP) with anti-HA coupled magnetic beads. *Antibody heavy chain; ** antibody light chain. **C.** Quantification of CTLA4-HA ubiquitylation relative to NT1 for data represented in (B). Ubiquitylated CTLA4-HA was normalised to total CTLA4-HA pulled down. Individual data points from two independent, colour-coded experiments are shown. Error bars indicate the range. **D.** Representative western blots of TUBES pulldown of ubiquitylated CTLA4. HeLa S3 Flp-In CTLA4-HA and A2058 cells were transfected as in (A and B) and cell lysates subjected to TUBES pulldown. **E.** Quantification of CTLA4-HA ubiquitylation relative to NT1 for data represented in (D). Ubiquitylated CTLA4(-HA) was normalised to total ubiquitin pulled down. Individual data points from two independent, colour-coded experiments are shown. Error bars indicate the range. **F.** Representative western blots of CHX chase in HeLa S3 Flp-In CTLA4-HA and A2058 cells following transfection with non-targeting (NT1) or HD-PTP siRNA. Cells were treated with CHX for indicated times before lysis. The half-life of CTLA4 was estimated using an exponential decay model. Error bars indicate standard deviation from three independent experiments.

The multi-domain structure of USP8 and intricate low-affinity interaction network ensures assembly of the many ESCRT-machinery components through co-incidence detection (Row et al., 2007). It is conceivable that it is the loss of USP8 as a scaffold that is relevant for efficient CTLA4 sorting to the lysosome, as opposed to its catalytic DUB function. In order to formally establish whether DUB activity of USP8 is essential for CTLA4 trafficking and degradation, we conducted a series of rescue experiments using siRNA-resistant GFP-tagged USP8 constructs. These were aimed at correcting CTLA4 turnover and endolysosomal accumulation, which can be visually assessed (Figure **7A, B**). As well as a catalytically inactive mutant (USP8 C786S), we also included a mutant with a deletion of the MIT domain (ΔMIT), which we previously showed is required for CHMP interaction and recruitment to endosomes (Row et al., 2007). Our results show that only wild-type, catalytically active and endosome-associated USP8 is able to restore CTLA4 down-regulation (Figure **7C, D**). In parallel, we monitored CTLA4 ubiquitylation status, which was likewise rescued by wild-type but not catalytically inactive or MIT-deleted mutants of USP8 (Figure **7E, F**).

**Figure 7.**
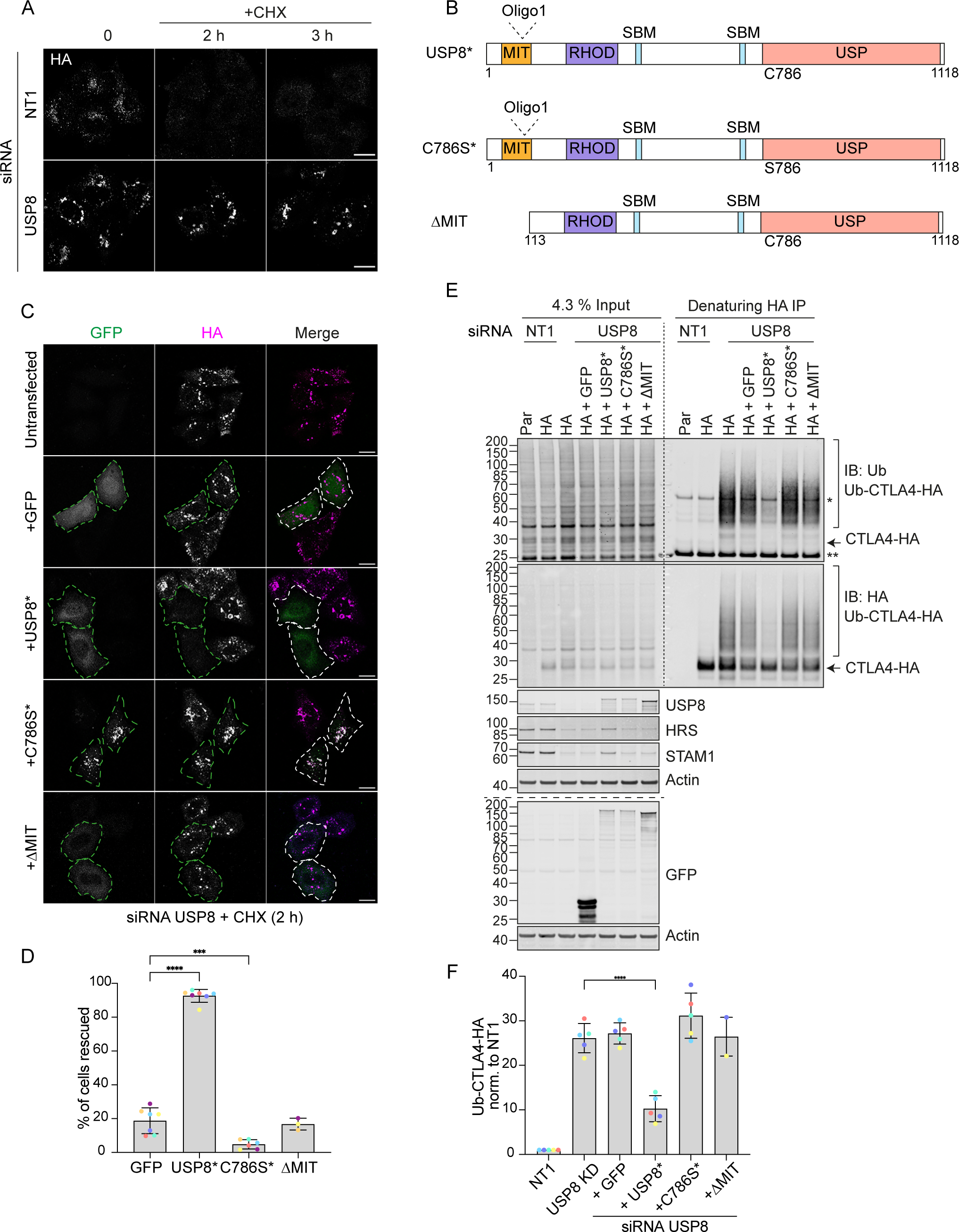
Catalytic activity and endosomal localisation of USP8 are essential for CTLA4 degradation and ubiquitylation. **A.** Representative confocal images of HeLa S3 Flp-In CTLA4-HA transfected for 72 h with NT1 or USP8 siRNAs before Cycloheximide (CHX) treatment for indicated times. Cells were fixed and stained for HA. Scale bar = 15 µm. **B.** GFP-tagged USP8 siRNA-resistant (USP8*) constructs used in this study. MIT, Microtubule interacting; SBD: SH3 domain binding motif; RHOD, Rhodanese homology domain; USP, Ubiquitin specific protease - catalytic domain. **C.** Representative confocal images of HeLa S3 Flp-In CTLA4-HA cells transfected with non-targeting (NT1) or USP8 siRNA and GFP, siRNA-resistant GFP-tagged USP8 (USP8*), catalytically inactive USP8 (C786S*) or ΔMIT-USP8. Cells were treated for 2 h with CHX prior to fixation and staining for HA. Scale bar = 15 µm. **D.** Quantification of cells showing rescued phenotypes calculated for data represented in (C). Individual data points from 3 (ΔMIT), 6 (C786S) or 7 (GFP and USP8*) independent, colour-coded experiments are shown. Error bars indicate the standard deviation. Total number of cells analysed: GFP (599); USP8* (666); C786S (486); ΔMIT (322). One-way ANOVA and Dunnett’s multiple comparison test, ***p<0.001 and ****p<0.0001. **E.** Representative western blots of ubiquitylated CTLA4-HA immunoprecipitated under denaturing conditions. HeLa S3 Flp-In parental (Par) or CTLA4-HA (HA) cells were transfected with NT1 or USP8 siRNA and either GFP or GFP-tagged siRNA-resistant USP8 constructs as in (C). Cells were lysed in denaturing SDS lysis buffer and lysates subjected to immunoprecipitation (IP) with anti-HA coupled magnetic beads. *Antibody heavy chain; ** antibody light chain. IB: Immunoblot. **F.** Quantification of CTLA4-HA ubiquitylation relative to NT1 for data represented in (E). Ubiquitylated CTLA4-HA was normalised to total CTLA4-HA pulled down. Individual data points from 2 (ΔMIT) or 5 independent, colour-coded experiments are shown. Error bars indicate the standard deviation (n=5) or range (n=2). One-way ANOVA and Dunnett’s multiple comparison test ,****p<0.0001.

### Ubiquitylation at Lys 203 and 213 promotes CTLA4 degradation

CTLA4 harbours 5 lysines in its cytoplasmic tail (Figure **8A**). We set out to identify which of these are critical for its ubiquitylation by generating a series of single and double point mutants as well as a “K-null” mutant in which all 5 lysines are converted to arginine. Deleting all lysines eliminated CTLA4 ubiquitylation, whilst mutating the three amino acids most proximal to the transmembrane domain had only negligible effects (Figure **8B, C**). Mutation of the last two lysines in the C-terminal tail together (K203R, K213R), all but abolished ubiquitylation, whilst each individual mutation reduced the signal by half. Importantly, mutation of these same lysines dramatically increases the stability of CTLA4 to a similar degree as mutation of all lysines (K-null) or ConcA treatment (Figure **8D,E** and **S4A-D**). In line with the slower turnover, the steady state distribution of both K-null and K203R/K213R mutants is partially shifted from late (LAMP1-positive) to early (EEA1-positive) endosomal compartments (Figure **S4E-H**).

**Figure 8.**
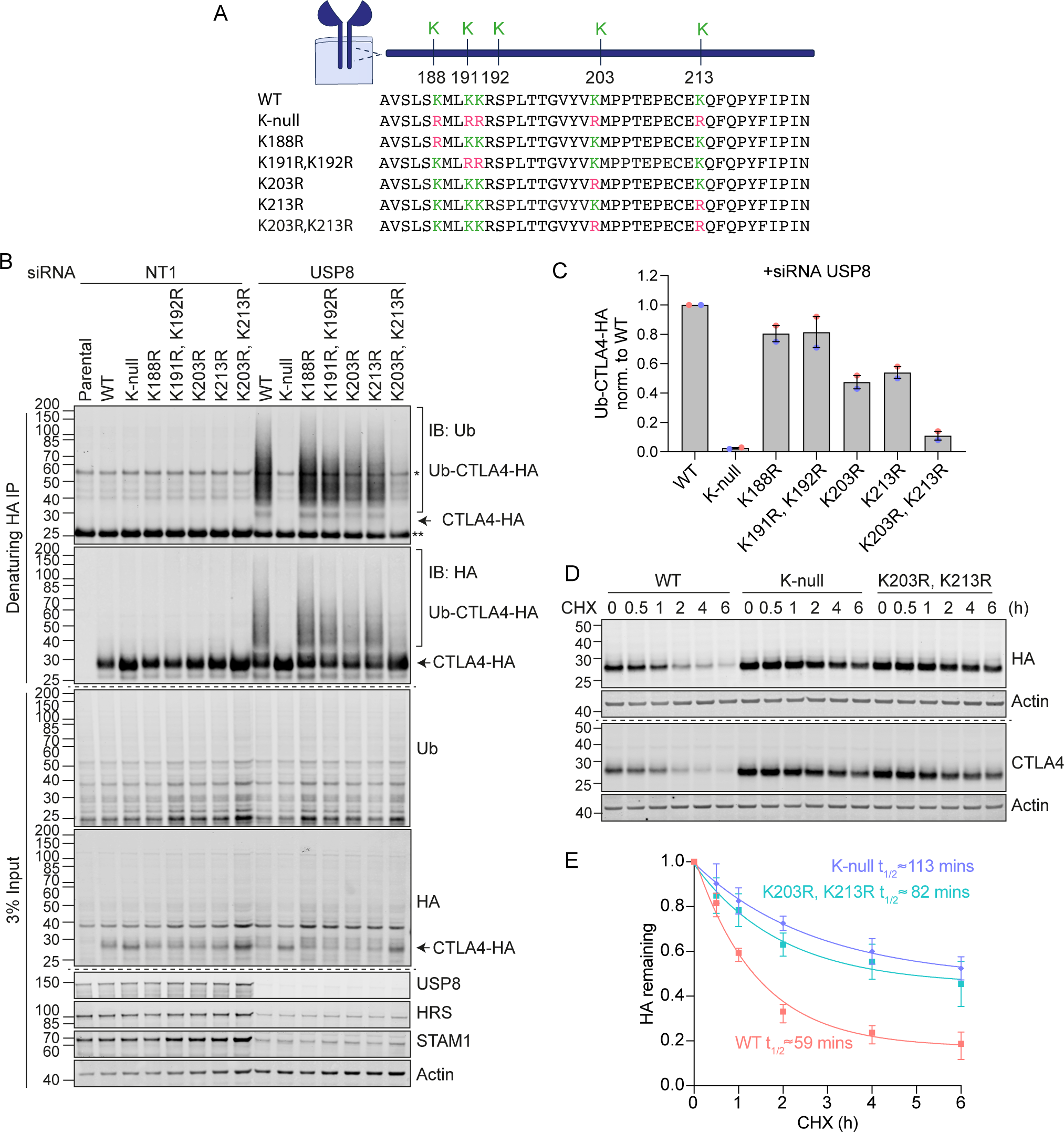
Ubiquitylation of CTLA4 on Lys203 and Lys213 is responsible for its rapid turnover. **A.** Depiction of the cytoplasmic tail of CTLA4 and the lysine mutants analysed in this study. **B.** Representative western blots of ubiquitylated CTLA4-HA immunoprecipitated under denaturing conditions. HeLa S3 Flp-In parental (Par) or CTLA4-HA WT or lysine mutant cells were transfected for 72 h with with non-targeting (NT1) or USP8 siRNA. Cells were lysed in denaturing SDS lysis buffer and lysates subjected to immunoprecipitation (IP) with anti-HA coupled magnetic beads. *Antibody heavy chain; ** antibody light chain. IB: Immunoblot. **C.** Quantification of ubiquitylated lysine mutant CTLA4-HA relative to WT for data represented in (B). Ubiquitylated CTLA4-HA is shown normalised to immunoprecipitated CTLA4-HA. Individual data points from two independent, colour-coded experiments are shown. Error bars indicate the range. **D.** Representative western blots of HeLa S3 Flp-In CTLA4-HA WT, K-null and K203R,K213R double lysine mutant cells treated with CHX for indicated times before lysis. **G.** Quantification of CTLA4-HA turnover for data represented in (D). The half-life of CTLA4-HA was estimated using an exponential decay model. Error bars indicate standard deviation from three independent experiments.

### UbiCRest analysis of CTLA4-associated polyubiquitin chains

We next set out to identify the ubiquitin chain linkages associated with CTLA4-HA by performing a ubiquitin chain restriction (UbiCRest) analysis. This assay leverages the chain linkage specificity of various DUBs to infer the linkage composition of ubiquitylated species associated with a protein of interest (Hospenthal et al., 2015). USP2 and vOTU show high promiscuity towards ubiquitin chain linkages and will also remove any proximal ubiquitin moieties (Ub-P). In addition to a high molecular weight smear, a prominent discrete band at ∼40kD, consistent with a dual mono-ubiquitylated form (Lys203, Lys213), was lost upon treatment with either enzyme (Figure **9A, B**). Of all the linkage-specific DUBs tested, only AMSH*, TRABID and OTUD2 removed ubiquitin chains off CTLA4-HA, which is best appreciated by focusing on the smear of protein above 60 kDa and the release of ubiquitin species into the supernatant (Figure **9A-D**). The ubiquitylation signal was clearly reduced upon treatment with the stringent Lys63 chain-directed enzyme AMSH* (Komander et al., 2009; McCullough et al., 2006). Direct conjugation of Lys63-linked chains with CTLA4 was also confirmed following cell lysis under denaturing conditions (Figure **9E**). OTUD2 was the most effective DUB for CTLA4-HA deubiquitylation but in distinction to AMSH* only effected partial loss of Lys63-linked ubiquitylation (Figure **9A**). Whilst Lys63 linkages have been prominently linked to endosomal trafficking, we reasoned that other linkage types must be present on CTLA4-HA (Clague et al., 2012). TRABID and OTUD2, which also reduced CTLA4 ubiquitylation, can both hydrolyse Lys29 and Lys33 ubiquitin chains, but only OTUD2 can remove Lys27 ubiquitin chains (Licchesi et al., 2012; Mevissen et al., 2013; Michel et al., 2015). More processing is evident when comparing OTUD2 with TRABID indicating the likely presence of Lys27 chains. We could confirm this by using a Lys27 ubiquitin-selective antibody, which detects Lys27-linked ubiquitylation on CTLA4-HA immunoprecipitated from denatured lysates (Figure **9F**). Accordingly, this Lys27 signal was untouched by either AMSH* or TRABID but reduced significantly by OTUD2 (Figure **9G**). We next analysed the ubiquitin that is released in the UbiCRest assay into the supernatant (from CTLA4 and associated proteins that specifically co-immunoprecipitate). We could see that this is incompletely processed, leaving residual oligomers indicative of heterotypic and possibly branched chain ubiquitin species (Figure **9A** **and S5**). For example, treatment with OTUD2 generates a higher molecular weight species, which is digested by co-incubation with AMSH* (Figure **S5**, K63-species). The composite picture that emerges is a complex mixture of mono-ubiquitin and Lys63 alongside more unusual Lys27 and Lys29 chain types.

**Figure 9.**
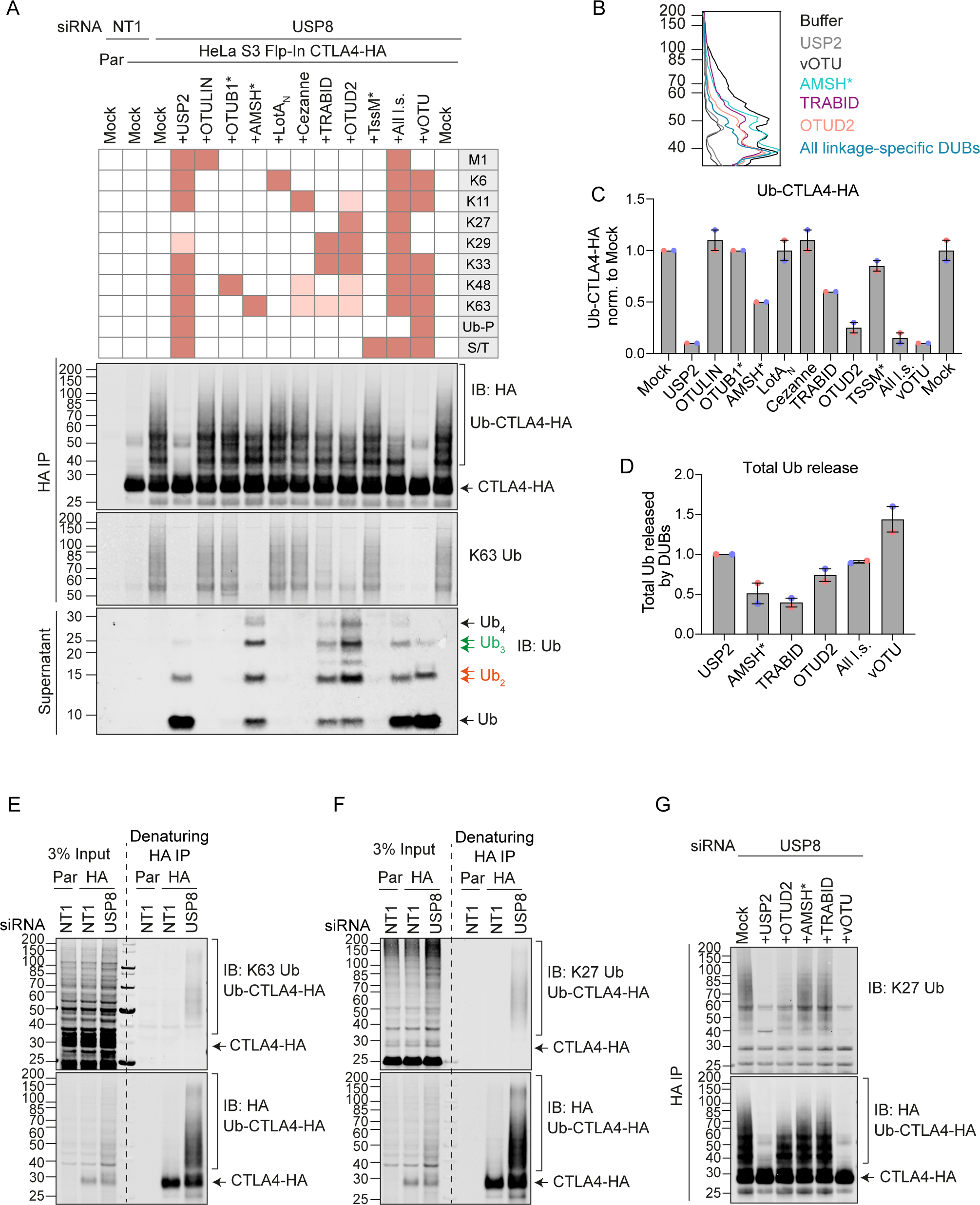
Ubiquitin Chain Restriction (UbiCRest) analysis reveals K63, K27 and K29 ubiquitin chain association with CTLA4-HA. **A.** Representative UbiCRest analysis western blots for CTLA4-HA. HeLa S3 Flp-In CTLA4-HA were transfected for 72 h with non-targeting (NT1) or USP8 siRNA prior to lysis and HA-immunoprecipitation (IP; IB: Immunoblot). CTLA4-HA-beads were treated with indicated DUBs or buffer only (Mock) for 1 h at 37°C, and analysed alongside supernatants (released Ubiquitin species). Specific activities of DUBs as reported in the literature are depicted in the key above the blot. Ub-P indicates ability to cleave proximal Ubiquitin; S/T denotes activity for serine or threonine linkages. All l.s.: All linkage specific DUBs (OTULIN, OTUB1*, AMSH*, LotA_N_, Cezanne, TRABID, OTUD2, TssM*). **B.** Line graphs corresponding to the Ub-CTLA4-HA signal shown in (A). **C.** Quantification of Ub-CTLA4-HA signal remaining after DUB-treatment shown relative to Mock (“buffer only” control) for data represented in (A). Individual data points from two independent, colour-coded experiments are shown. Error bars indicate the range. **D.** Quantification of total ubiquitin released into the supernatant by DUBs relative to USP2 for data represented in (A). Individual data points from two independent, colour-coded experiments are shown. Error bars indicate the range. **E, F.** Representative western blots of CTLA4-HA immunoprecipitated (IP) under denaturing conditions probed with K63 (E) and K27 (F) ubiquitin chain linkage-specific antibodies. **G.** Representative western blot showing K27-linked ubiquitin associated with immunoisolated CTLA4-HA after UbiCRest treatment with indicated DUBs.

## Discussion

There is a mismatch between the prominence of CTLA4 within the immuno-therapeutic space and our rudimentary understanding of its dynamics and trafficking itinerary. Pioneering work from the Sansom laboratory has shown that exogenous expression of CTLA4 in epithelial-derived cells can broadly capture its behaviour in T cells (Janman et al., 2021; Qureshi et al., 2012; Rowshanravan et al., 2018). Here we have adopted a similar approach of exogenous expression, but using a Flp-In cell line, to create isogenic cell panels that allow for highly accurate comparison between mutants. In parallel, we have also utilised melanoma and squamous lung carcinoma cell lines that naturally express CTLA4. In all cases cycloheximide chase experiments confirm the short half-life previously reported, which is significantly less than one hour in our hands (Qureshi et al., 2012). This is highly unusual, as only ∼5% of the proteome has a half-life less than 8 hours (Li et al., 2021; Rusilowicz-Jones et al., 2022).

The principal degradation pathways are normally discriminated by application of v-ATPase inhibitors (for lysosomal degradation) or proteasome inhibitors (Rusilowicz-Jones et al., 2022). Here we show that the major CTLA4 turnover pathway is lysosomal, with no evidence for a proteasomal contribution despite the rapid turnover. There remains scope for minor alternate pathways, including secretion in the form of exosomes demonstrated herein. Although ubiquitylation of CTLA4 has been previously noted, its significance has not been directly tested (Kennedy et al., 2022). Here, using acute application of the ubiquitin E1 enzyme inhibitor TAK-243, we show that active ubiquitylation is absolutely required for CTLA4 degradation. Within the cytoplasmic tail of CTLA4, we have been able to show that direct ubiquitylation of just two out of the five candidate Lys residues governs stability. This opens up a new vista for identifying druggable targets within the ubiquitin system which might regulate CTLA4 expression. For example regulatory DUBs have been established as tractable targets for selective small molecule inhibition (Clancy et al., 2021; Harrigan et al., 2018; Turnbull et al., 2017).

At this point we have taken a candidate-based approach, focusing on the two DUB enzymes which are most prominent at endosomes, the Lys63-chain specific metalloenzyme AMSH and the non-selective USP cysteine protease family member USP8 (Clague et al., 2019; Urbe et al., 2012). Lys63-linked ubiquitin chain modifications have been shown to be critical for efficient lysosome-directed sorting of multiple receptor types, although the effects of AMSH loss tend to be modest (Barriere et al., 2007; Clague et al., 2012). We find no effect of AMSH depletion on CTLA4 stability. In contrast, USP8 depletion enhances both CTLA4 half-life and ubiquitylation. If ubiquitylation is the critical signal for lysosomal sorting, one might expect this to lead to enhanced degradation, but the opposite holds true. This recalls past findings of MET, EGFR and CXCR4 receptor degradation which are likewise delayed by USP8 depletion (Berlin et al., 2010; Bowers et al., 2006; Row et al., 2006; Savio et al., 2016). The effects of USP8 loss on endosomes are highly pleiotropic and include the clustering of endosomes as well as the loss of the ESCRT-0 components and ubiquitin binding proteins, HRS and STAM (Clague and Urbe, 2017). The block to degradation of CTLA4 cannot be accounted for by loss of ESCRT-0 alone, as depletion of HRS has no effect. Two lines of argument show that USP8 must associate with endosomes to exert this effect. Depletion of a key adapter protein critical for USP8 localisation, HD-PTP, phenocopies the block to CTLA4. Secondly, deletion of the MIT domain from siRNA-resistant USP8, which is required for localisation, renders it incapable of rescuing the depleted phenotype.

We were particularly keen to understand if this regulation by USP8 is maintained in T cells. CTLA4 is constitutively expressed by FoxP3^+^ regulatory T cells cells (T_Regs_) and up-regulated upon activation in conventional T cells. Mice with a T cell-specific inactivation of USP8 (Usp8f/fCd4Cre) develop colitis, which is mediated by CD8^+^ γδT cells in concert with dysfunctional T_Regs_ (Dufner et al., 2015). Colitis is also among the most frequent and problematic immune-mediated adverse events that are associated with dual checkpoint inhibition and a combination of anti-CTLA4 and anti-PD-1 monoclonal antibodies has been shown to exacerbate dextran sulphate sodium-induced autoimmune colitis in mice (Perez-Ruiz et al., 2019; Postow et al., 2018). We speculate that the disequilibrium imposed on CTLA4 following USP8 deletion may contribute to the colitis phenotype.

The lack of impact of AMSH upon CTLA4 may reflect the involvement of chain types other than its exclusive substrate, Lys63-linked chains, that have been most prominently linked to endocytosis (Clague et al., 2012). We believe this is why USP8 but not AMSH regulates ESCRT-0 stability, despite their sharing of many interactions (Clague et al., 2012; Clague and Urbe, 2006). AMSH will not cleave a proximal ubiquitin and we here provide evidence for a significant population of dual mono-ubiquitylated CTLA4, which provides a relatively weak endosomal sorting signal for other receptor types (Haglund et al., 2003; Ma et al., 2012). Our UbiCRest chain analysis comes with the caveat that we are analysing modifications which accrue in the absence of USP8, for otherwise the ubiquitylation is too labile for such analysis. As USP8 is a relatively promiscuous enzyme for different chain linkages, the most parsimonious interpretation is that the palette we reveal reflects that which may be occurring in unperturbed cells. If anything, a slight bias against Lys27 and Lys29 linkages is predicted because these are not particularly good USP8 substrates, at least in their diubiquitin form (Ritorto et al., 2014; van Tol et al., 2023). We provide evidence for a heterogeneous, partially branched, population that includes poorly characterised linkages not previously directly associated with receptor trafficking. However, two ESCRT machinery-linked proteins have been shown to have Lys27/29 (STAM) or Lys29/48 (VPS34) ubiquitin modifications (Chen et al., 2021; McElrath et al., 2023). The Parkinson’s disease associated kinase LRRK2 that has a role in endolysosomal stress responses can also be regulated by K27 and K29-linked ubiquitin chain modifications (Nucifora et al., 2016).

Heterotypic branched chains have recently been shown to enhance the rate of proteasomal degradation of cytosolic substrates (Meyer and Rape, 2014). We propose that a similar principle holds true for lysosomal sorting of short-lived membrane proteins such as CTLA4, which we show presents a complex branched ubiquitin chain profile composed of Lys63, Lys27 and Lys29 linkages. The present study opens up the ubiquitin system as a key player in CTLA4 biology that likely includes specific druggable regulators. So far biological agents such as antibodies have been used to effect changes in CTLA4 function for clinical purposes. We envision that future developments of small molecule regulators of CTLA4 stability may provide a complementary approach.

## Materials and Methods

### Cell culture

A2058 cells were cultured in DMEM supplemented with GlutaMAX and 10% FBS; HeLa S3 Flp-In in DMEM supplemented with GlutaMAX, 10% FBS and 1x non-essential amino acid (NEAA); NCI-H520 in RPMI supplemented with GlutaMAX and 10% FBS at 37°C and 5% CO2. Cells were routinely screened for mycoplasma infection.

### Generation of HeLa S3 Flp-In CTLA4-HA and CTLA4-HA lysine mutants

Human CTLA4-HA flanked by BglII/EcoRI sites was synthesised and provided in pTWIST-High-Kan by TWIST Bioscience, USA. CTLA4-HA was sub-cloned into pcDNA3.1 using its BamHI/EcoRI sites and then subcloned into a pEF5/FRT plasmid using KpnI/EcoRV restriction sites. For the generation of the CTLA4-HA lysine mutants (K-null; K188R; K191R/K192R; K203R; K213R; K203R/K213R), gene fragments flanked by BamHI/NotI sites, encoding the cytoplasmic tail of CTLA4 encompassing the relevant mutations (132 to 223) and a C-terminal HA tag, were synthesised by TWIST, sub-cloned into pEF5/FRT/CTLA4-HA and sequence-verified. To generate stable cell lines expressing wild-type and CTLA4-HA lysine mutants, HeLa S3 Flp-In host cells were co-transfected with pEF5/FRT/CTLA4-HA and pOG44 at a ratio of 1:9 using GeneJuice (Merck Millipore, 70967) according to manufacturer’s instructions. Transfected cells were selected using Hygromycin B (Invitrogen, 10687010, 150 µg/ml). For WT CTLA4, individual clones were amplified and the lysine mutant expressing cells were maintained as pools.

### Mouse T cell isolation, expansion and USP8 deletion

T cells were isolated from spleens of sex-matched 8-12-week-old Rosa26-CreERT2 (Gt(ROSA)26Sortm2(cre/ERT2)Brn) USP8 fl/fl mice and USP8fl/fl (USP8^tm1floxedKPK^) littermate control (Hameyer et al., 2007; Niendorf et al., 2007) using BD IMag^TM^ Biotinylated Mouse CD4 T lymphocyte enrichment Cocktail and BD IMAG^TM^ Streptavidin Particles Plus according to manufacturer’s instructions. *Ex vivo* expansion of T cells was performed using anti-CD3 (eBioscience, clone 145-2C11, 1 µg/ml) and anti-CD28 antibodies (eBioscience, clone 37.51, 1 µg/ml) on TC plates pre-coated with anti-hamster antibody (BD Pharmingen, G94-56), and IL-2 (Immunotools, 100 U/ml) for continued cell culture. 4-hydroxytamoxifen (OHT, Sigma, 1 µM) was added 24 h post T cell isolation and stimulation. Cells were lysed 72 h post OHT addition.

### Transfection and siRNA interference

Cells were treated with 40 nM non-targeting (NT1) or target-specific siRNA oligonucleotides (Dharmacon) using Lipofectamine RNAi-MAX (Invitrogen, 13778030) according to manufacturer’s instructions. The medium was exchanged 24 h after transfection and cells were harvested 72 h post-transfection. For plasmid transfection, GeneJuice (Merck Millipore, 70967) was used according to manufacturer’s instruction. For rescue experiments, endogenous USP8 was depleted by transfecting cells with USP8 oligo 1 (40 nM) using RNAiMAX for 72 hours, followed by GFP or GFP-USP8 siRNA-resistant constructs using GeneJuice for the last 46 h prior to lysis or fixation.

### siRNA and plasmids

All siRNA oligonucleotides were obtained from Dharmacon, Horizon Discovery: CTLA4 (SMARTPool, L-016267-00-0005: 5’-GAAGCCCUCUUACAACAGG-3’, 5’- GAACCCAGAUUUAUGUAAU-3’, 5’-GUAUGCAUCUCCAGGCAAA-3’, 5’- GGACUGAGGGCCAUGGACA-3’), USP8 oligo 1 (siGNEOME, D-005203-02: 5’- UGAAAUACGUGACUGUUUAUU-3’), custom-made AMSH oligo 2 (CTM-931180: 5’- UUACAAAUCUGCUGUCAUUUU-3’), HD-PTP (SMARTPool, L-009417-00-0005: 5’- GUGCACAGGUGGUAGAUUA-3’, 5’-GCAAACAGCGGAUGAGCAA-3’, 5’- GCAUGAAGGUCUCCUGUAC-3’, 5’-GUAGUGUCCUCCGCAAGUA-3’), HRS (SMARTPool, L- 016835-00-0005: 5’-GAGGUAAACGUCCGUAACA-3’; 5’-GCACGUCUUUCCAGAAUUC-3’; 5’- AAAGAACUGUGGCCAGACA-3’; 5’-GAACCCACACGUCGCCUUG-3’), Syntenin (SMARTPool, L- 008270-00-0005: 5’-GGAGAGAAGAUUACCAUGA-3’, 5’-GACCAAGUACUUCAGAUCA-3’, 5’- GGAUGGUCUUAGAAUAUUU-3’, 5’-GCAUUUGACUCUUAAGAUU-3’), ALIX (SMARTPool, L- 004233-00-0005: 5’-CAGAUCUGCUUGACAUUUA-3’; 5’-UCGAGACGCUCCUGAGAUA-3’; 5’- GCGUAUGGCCAGUAUAAUA-3’; 5’-GUACCUCAGUCUAUAUUGA) and non-targeting 1 (NT1) control (ON-TARGETplus, D-001810-01, 5’-UGGUUUACAUGUCGACUAA-3’).

The generation of GFP-USP8* and GFP-ΔMIT-USP8 have been described previously (Row et al., 2006; Row et al., 2007). GFP-USP8*-C786S was generated by performing QuickChange site-directed mutagenesis using GFP-USP8* as the template and the following primer pairs: 5’-ACTTAGGAAATACTAGTTATATGAACTCA-3 ’and 3’-TGAGTTCATATAACTAGTATTTCCTAAGT-5’. Transformants were screened by restriction digestion and validated by sequencing.

### Antibodies and reagents

Antibodies and other reagents used were as follows: anti-human CTLA4 E1V6T (Cell Signalling Technology, 96399, WB 1:500), anti-mouse CTLA4 (R&D Systems, AF476, WB 1:1000), anti-human CTLA4 BNI3 (BD Pharmingen, 555851, IF 1:100), anti-CTLA4 F8 (Santa Cruz Biotechnology, sc-376016, WB 1:500), anti-USP8 (R&D Systems, AF7735, WB 1:500), anti-USP8 (Bethyl, A301-350A, WB 1:2000), anti-mouse USP8 X39 (a gift from E. Martegani, WB 1:2000), anti-AMSH (Homemade, rabbit 850, WB 1:1000), anti-HRS (In house, rabbit 864/3, WB 1:1000), anti-HRS (Everest Biotech, EB07211, WB 1:2000), anti-HRS (abcam, ab155539, WB 1:1000, IF 1:200), anti-STAM1 (abcam, ab155527, WB 1:1000), anti-ubiquitin VU1 (LifeSensor, VU101, WB 1:2000; Fig1C, Fig1E), anti-ubiquitin U5379 (Sigma-Aldrich, U5379, WB 1:1000; Fig9A, FigS5), anti-ubiquitin FK2 (Enzo, PW8810, WB 1:1000, Fig4A, Fig4C, Fig4E, Fig6B, Fig6D, Fig7E, Fig8B), anti-ubiquitin Lys63-specific clone Apu3 (Millipore, 05-1308, WB 1:1000), anti-ubiquitin Lys27-specific (Abcam, 181537, WB 1:1000), anti-HDPTP (ProteinTech, 10472-1-AP, WB 1:500), anti-ALIX (Santa Cruz Biotechnology, sc-53540, WB 1:1000), anti-Syntenin 1 (Novus Biologicals, H00006386-B01P, WB 1:1000), anti-TOMM20 (BD Transduction, 612278, WB 1:1000), anti-CD63 (Novus Biologicals, NBP2-42225SS, WB 1:500), anti-HA (Novus Biologicals, NB600-362, WB 1:10000, IF 1:250), sheep anti-GFP (gift from Ian Prior, University of Liverpool, WB 1:1000), anti-Actin (Proteintech, 66009, WB 1:10000), anti-Actin (Sigma, A2066, WB 1:10000), anti-γ-Tubulin (Abcam, ab11317, WB 1:1000), anti-LAMP1 D2D11 (Cell Signalling Technology, 9091, IF 1:400), anti-EEA1 (BD Transduction Laboratories, 610456, IF 1:500), anti-EEA1 (In house, rabbit 243/3, IF 1:1000); Cycloheximide (Sigma-Aldrich, C7698, 100 µg/ml), Concanamycin A (Sigma-Aldrich, C9705, 100 nM), Folimycin A (Sigma-Aldrich, 344085, 100 nM), Epoxomicin (Millipore, 324800, 100 nM), TAK-243 (Selleckchem, S8341, 1 µM).

### Cell lysis and western blot analysis

Cultured cells were washed twice in ice-cold PBS and lysed in NP40 lysis buffer (0.5 % NP40, 25 mM Tris pH 7.5, 100 mM NaCl and 50 mM NaF) supplemented with mammalian protease inhibitor cocktail (Sigma-Aldrich, P8340) and PhosSTOP (Roche, 49068450001) for 10 mins on ice. Lysates were clarified by centrifugation and protein concentration determined using the Pierce BCA protein assay according to manufacturer’s instructions. Samples were diluted with 5 x SDS-sample buffer (15 % w/v SDS, 312.5 mM Tris-HCl pH 6.8, 50 % glycerol and 16 % beta-mercaptoethanol) and boiled at 95°C. Proteins were resolved using SDS–PAGE (Invitrogen NuPage gel 4–12 %), transferred to nitrocellulose membrane (10600001 or 10600002; Amersham Protran 0.2 and 0.45 µm pore size), stained with Ponceau S staining solution (P7170; Sigma-Aldrich), blocked in 5 % milk (Marvel) or 0.5% fish skin gelatin (Sigma, G7765) in TBST (TBS: 20 mM Tris–Cl, pH 7.6 and 150 mM NaCl, supplemented with Tween-20 (10485733; Thermo Fisher Scientific)) before incubation with primary antibodies overnight. Visualisation and quantification of western blots were performed using IRdye 800CW (anti-mouse 926-32212, anti-rabbit 926-32213 and anti-goat 926-32214) and IRdye 680LT (anti-mouse 926-68022, anti-rabbit 92668023 and anti-goat 926-68024) coupled secondary antibodies and an Odyssey infrared scanner (LI-COR Biosciences). For western blot quantification, raw signal values were obtained using ImageStudio Lite (Li-COR) following background subtraction and the raw values of each condition were normalised to the average of the quantified raw values from each individual blots. For stripping and reprobing, membranes were incubated in 2 % SDS, 62.5 mM Tris-HCl pH 6.7 and 100 mM β-mercaptoethanol for 30 mins in a 50°C water bath, then washed three times in 0.2 % TX100 in PBS, followed by incubation in blocking buffer.

### Immunoprecipitation

For co-immunoprecipitation experiments, HeLa S3 Flp-In CTLA4-HA cells were lysed in NP40 lysis buffer as described above. Clarified lysates were incubated with 25 µl of anti-HA magnetic beads (Thermo Scientific, 88837) overnight at 4°C. Beads were washed three times with 0.05 % TBST and proteins eluted in sample buffer. For denaturing immunoprecipitation of CTLA4-HA, cells were washed twice with pre-warmed PBS and lysed in denaturing SDS lysis buffer (2% w/v SDS, 1 mM EDTA and 50 mM NaF) at 110°C, followed by boiling for 10 mins with intermittent vortexing. Lysates were diluted with 4 volumes of dilution buffer (2.5 % Triton X-100, 12.5 mM Tris pH 7.5 and 187.5 mM NaCl) before incubation with 25 µl of anti-HA magnetic beads (Thermo Scientific, 88837) at 4°C overnight. Beads were washed with TX100-SDS wash buffer (2 % Triton X-100, 0.4 % SDS, 10 mM Tris pH 7.5, 150 mM NaCl) and proteins were eluted in 50 mM NaOH. 1M Tris pH 8.5 was added to the eluted samples to neutralise the pH and samples were prepared in 5 x “hot lysis” sample buffer (7 % (w/v) SDS, 312.5 mM Tris-HCl pH 6.8, 50 % (w/v) glycerol and 16 % ß-mercaptoethanol).

### Tandem Ubiquitin Binding Entities (TUBES) pulldown

Cells were washed twice with ice-cold PBS and lysed in TUBES lysis buffer (50 mM Tris-HCl pH 7.5, 150 mM NaCl, 1 mM EDTA, 1 % (w/v ) NP40, 10 % (w/v) glycerol) supplemented with mammalian protease inhibitor cocktail (Sigma-Aldrich, P8340), PhosSTOP (Roche, 4906845001) and 10 mM NEM (Sigma-Aldrich, E3876-5G). Lysates were incubated with 20 µl (50% slurry) TUBES (Life sensors, UM402) or control agarose resin (Life sensors, UM400) overnight at 4°C. Beads were washed with 0.1 % TBST and proteins were eluted in sample buffer at 95°C.

### TCA-precipitation of proteins from media

Media were collected from 6-well plates and centrifuged at 3000 g for 15 mins to remove cell debris. SDS was added to the media at a final concentration of 0.02 %, followed by incubation on ice for 30 mins. Trichloroacetic acid (TCA, Sigma-Aldrich, T0699) was added to a final concentration of 10 % for a further 1 h incubation on ice, followed by centrifugation at 16,200 g for 30 mins at 4°C. The supernatant was removed and pellets were washed twice in ice-cold acetone. Pellets were air-dried and resuspended in SDS-sample buffer before analysis by SDS-PAGE and western blot.

### Exosome enrichment by serial centrifugation

All centrifugation steps were carried out at 4°C. Media were collected from 6-well plates and centrifuged sequentially at 300 g for 20 mins x 2 to remove cells, at 2000 g for 20 mins to remove large cell debris, at 10,000 g for 20 mins to remove smaller cell debris (using a Himac Ultracentrifuge and a S55A2 rotor). The supernatants were collected and centrifuged at 100,000 g for 70 mins using the same rotor to collect the exosomal fractions. The resulting pellets were washed in ice-cold PBS and re-pelleted again at 100,000 g for another 70 mins using the same rotor. The exosome-enriched pellets were resuspended alongside the 10,000 g pellets in SDS-sample buffer and analysed by SDS-PAGE and western blot.

### Proteinase K protection assay

To determine the topology of exosomal CTLA4, exosome-enriched pellets were resuspended in PBS containing 2.5 mM CaCl2 and 1 mM MgCl2 (PBS^++^). The samples were treated with proteinase K (Merck, P2308, 100 µg/ml) in 50 mM Tris-HCl pH 7.4 supplemented with 10 mM CaCl_2_ for 1 h at 37 °C in the absence or presence of 1 % (w/v) Triton X-100 in PBS^++^. To terminate the proteinase activity, phenylmethylsulfonyl fluoride (PMSF, Sigma-Aldrich, 78830, 2 mM final concentration) was added to the samples and incubated on ice for 5 mins. The samples were prepared in sample buffer as described above.

### Protein expression and purification

H. sapiens OTUD2 (aa 1-348; pOPIN-K), H. sapiens OTULIN (aa 1-352; pOPIN-B), H. sapiens Cezanne (aa 129-438; pOPIN-E), D. melanogaster TRABID (aa 318-778; pOPIN-S), Crimean Congo Hemorrhagic Fever Virus vOTU (aa 1-183; pOPIN-K), and L. pneumophila LotA_N_ (aa 1-300; pOPIN-B) were prepared as described previously (Akutsu et al., 2011; Keusekotten et al., 2013; Mevissen et al., 2013; Warren et al., 2023; Xia et al., 2022). OTUB1* and AMSH* are constitutively active engineered fusion proteins of H. sapiens UBE2D2(C85S)-OTUB1 and STAM-AMSH, expressed from the pOPIN-B vector as described previously (Michel et al., 2015). B. pseudomallei TssM* encodes the bacterial USP-type DUB engineered with a V466R mutation that enhances selectivity for ester-over isopeptide-linked ubiquitin, expressed from the pET-M30 vector as described (Szczesna et al., 2023). Briefly, all DUBs were expressed in E. coli Rosetta cells at 18 °C for 16 h following induction with 0.2-0.5 mM IPTG. Cells were harvested in 25 mM Tris (pH 7.4), 200 mM NaCl, 2 mM ß-mercaptoethanol (Buffer A), incubated with lysozyme and protease inhibitor cocktail (Millipore, Sigma), and lysed by sonication. Clarified lysate was applied to cobalt affinity resin (ThermoFisher), washed with additional Buffer A, and eluted with Buffer A containing 250 mM imidazole. Eluates were dialysed back into Buffer A overnight at 4 °C prior to concentration with 10 kDa MWCO Amicon centrifugal filters (Millipore, Sigma), quantification by absorbance at 280 nm, and flash freezing for storage at -80 °C. With the exception of OTUB1*, all His-tagged constructs expressed from pOPIN-B and pOPIN-E were left intact, while the His-GST and His-SUMO tags encoded by the pOPIN-K, pET-M30, and pOPIN-S constructs were cleaved by 3C, TEV, or SUMO proteases, respectively.

### Ubiquitin Chain Restriction (UbiCRest) assay

Cells were lysed in NP40 lysis buffer (0.5% NP40, 25 mM Tris pH 7.5, 100 mM NaCl and 50 mM NaF) supplemented with mammalian protease inhibitor cocktail (Sigma-Aldrich, P8340), PhosSTOP (Roche, 49068450001) and 20 mM NEM (Sigma-Aldrich, E3876-5G). The lysates were incubated with anti-HA magnetic beads (Thermo Scientific, 88837) overnight at 4°C to immunoprecipitate CTLA4-HA. The UbiCRest assay was performed in a 25 µl reaction volume at 37°C for 1 h in a thermoshaker at 900 rpm as previously described (Hospenthal et al., 2015). The DUBs used in this study are USP2 (Enzo, BML-UW9850-0100, 1 µM and Boston Biochem, E-504-0650, 1 µM), vOTU (1 µM), OTULIN (1 µM), OTUB1* (1 µM), AMSH* (1 µM), LotA_N_ (1 µM), Cezanne (0.2 µM), TRABID (0.2 µM), OTUD2 (1 µM) and TssM* (1 µM). The supernatants were collected after the incubation and prepared in sample buffer to analyse ubiquitin released during the UbiCRest assay. Proteins were eluted from the anti-HA magnetic beads at 60°C in sample buffer.

### Immunofluorescence

Cells were either fixed with 4 % paraformaldehyde (PFA, Agar Scientific AGR1026) in PBS or ice-cold methanol. Excess PFA was quenched with 50 mM NH_4_Cl/PBS and cells permeabilised with 0.2% Triton X-100 in PBS. All cells were incubated for 30 mins in blocking solution (3 % BSA in PBS), then stained with primary antibodies (1 h), followed by AlexaFluor-488- or AlexaFluor-594-coupled secondary antibodies (30 min) in blocking buffer. Coverslips were mounted onto glass slides using Mowiol containing DAPI. Fixed coverslips were imaged using a LSM900 Airyscan confocal microscope (63 x oil objective, acquisition software Zen Blue). All images were acquired sequentially and processed using Fiji (version 2.1.0) and Adobe Photoshop (version 24.5.0) software. Pearson’s and Mander’s coefficients (M2; fraction of CTLA4(-HA) colocalising with EEA1 or LAMP1 respectively) were measured using the JACoP plugin in Fiji (Bolte and Cordelieres, 2006).

### Statistical analysis

Graphs were plotted using GraphPad Prism10. Statistical significance was determined using one-way ANOVA (Figures 1D, 1F, 7D, 7F), unpaired t-test (Figures S2D, 5C-H) or two-way ANOVA with uncorrected Fisher’s LSD (Figure S4D). P-values are represented as *p<0.05 , **p<0.01, ***p<0.001 and ****p<0.0001.

## Acknowledgements

PYT was supported by a Wellcome Trust PhD studentship 102172/B/13/Z. MC is a Royal Society Industry Fellow INF\R2\212031. AD and KPK received funding from the Deutsche Forschungsgemeinschaft (DFG, German Research Foundation) under Germany’s Excellence Strategy (CIBSS–EXC-2189–Project ID 390939984) and DFG grant KN590/4-2. JNP is supported by the National Institute of General Medical Sciences (R35GM142486).

## Supplementary Figure Legends

**Figure S1.**
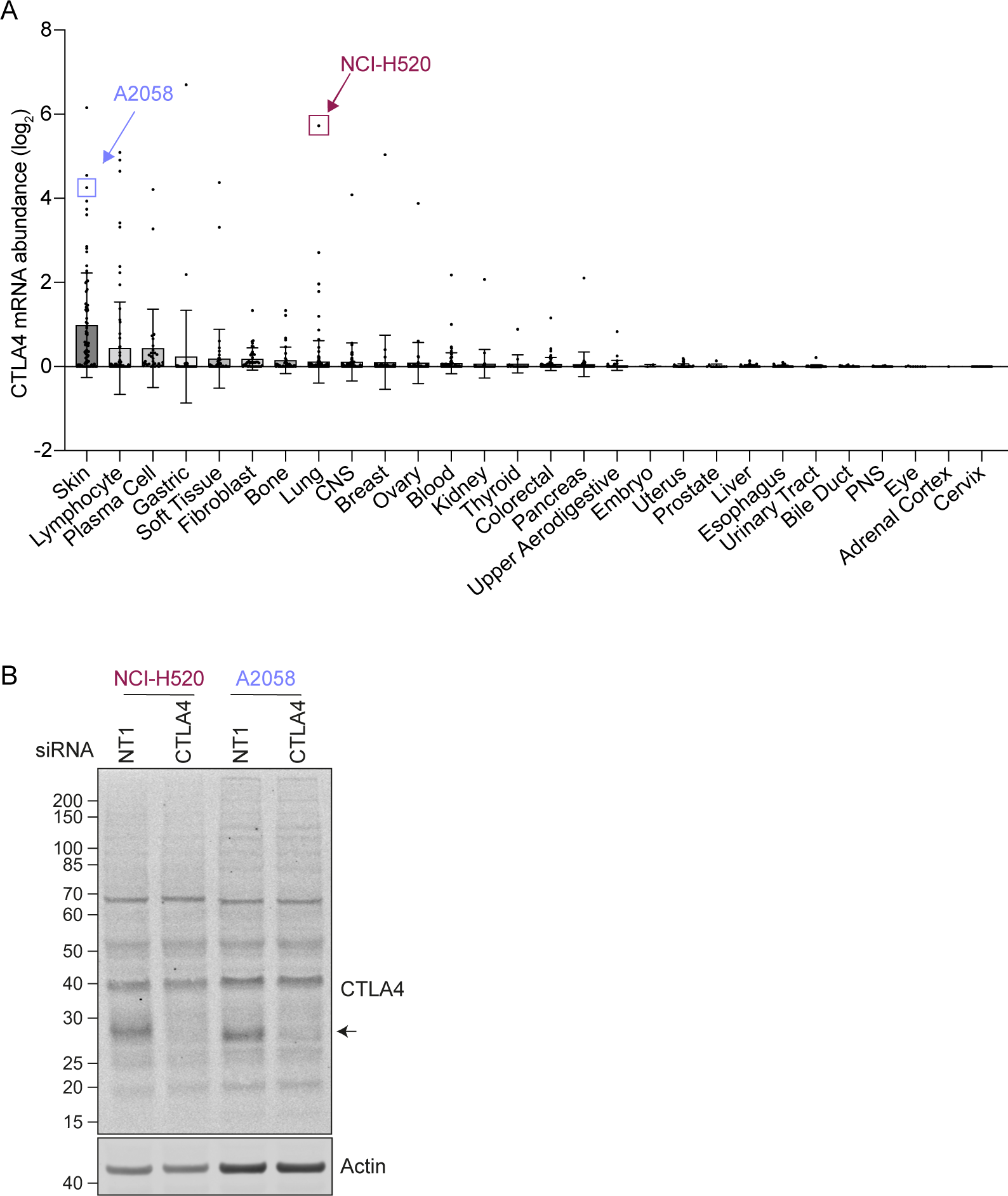
CTLA4 expression in cancer cell lines. **A.** mRNA levels derived from the Cancer Cell Line Encyclopaedia (CCLE) database (Barretina et al., 2012). Each dot represents an individual tumour cell line within the indicated categories. **B.** Representative western blot of NCI-H520 and A2058 cells transfected with siRNA against non-targeting (NT1) or CTLA4 for 72 h prior to lysis.

**Figure S2.**
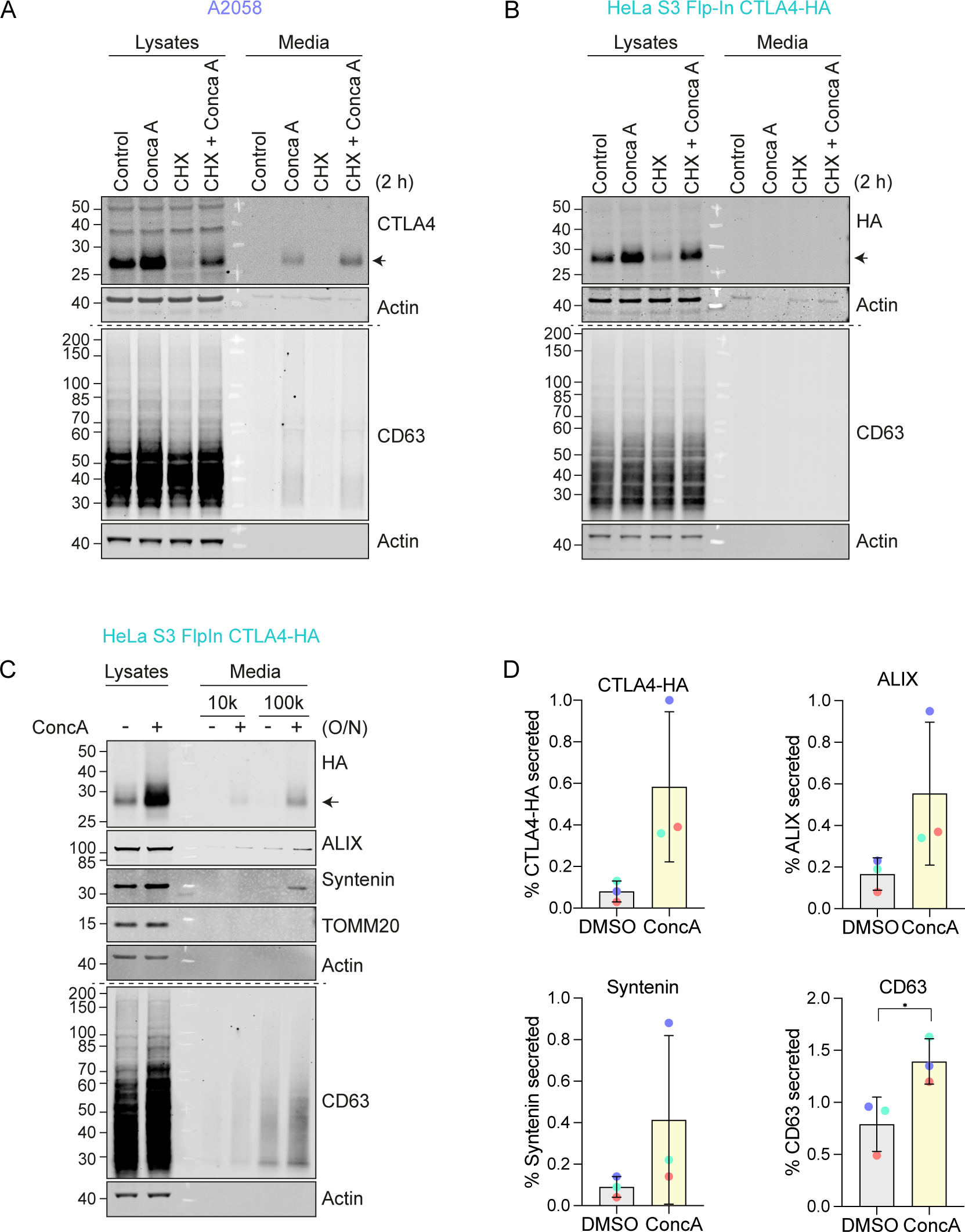
CTLA4 is secreted via exosomes from A3058 and HeLa S3 Flp-In CTLA4-HA cells. **A,B.** Representative western blots of the lysates and cultured supernatants from mock- and Concanamycin A (ConcA)-treated A2058 and HeLa S3 Flp-In CTLA4-HA cells. Cells were lysed and media subjected to trichloroacetic acid (TCA) precipitation. **C.** Representative western blots of the lysates and cultured supernatants from mock- and ConcA-treated HeLa S3 Flp-In CTLA4-HA cells for indicated times. Cells were lysed and the media collected by serial centrifugation to concentrate extracellular vesicles (100k pellet, exosome fraction). **D.** Quantification of CTLA4, Syntenin and CD63 secreted in the exosome fractions (100k) for data represented in (C). Individual data points from three independent, colour-coded experiments are shown. Error bars show standard deviation.

**Figure S3.**
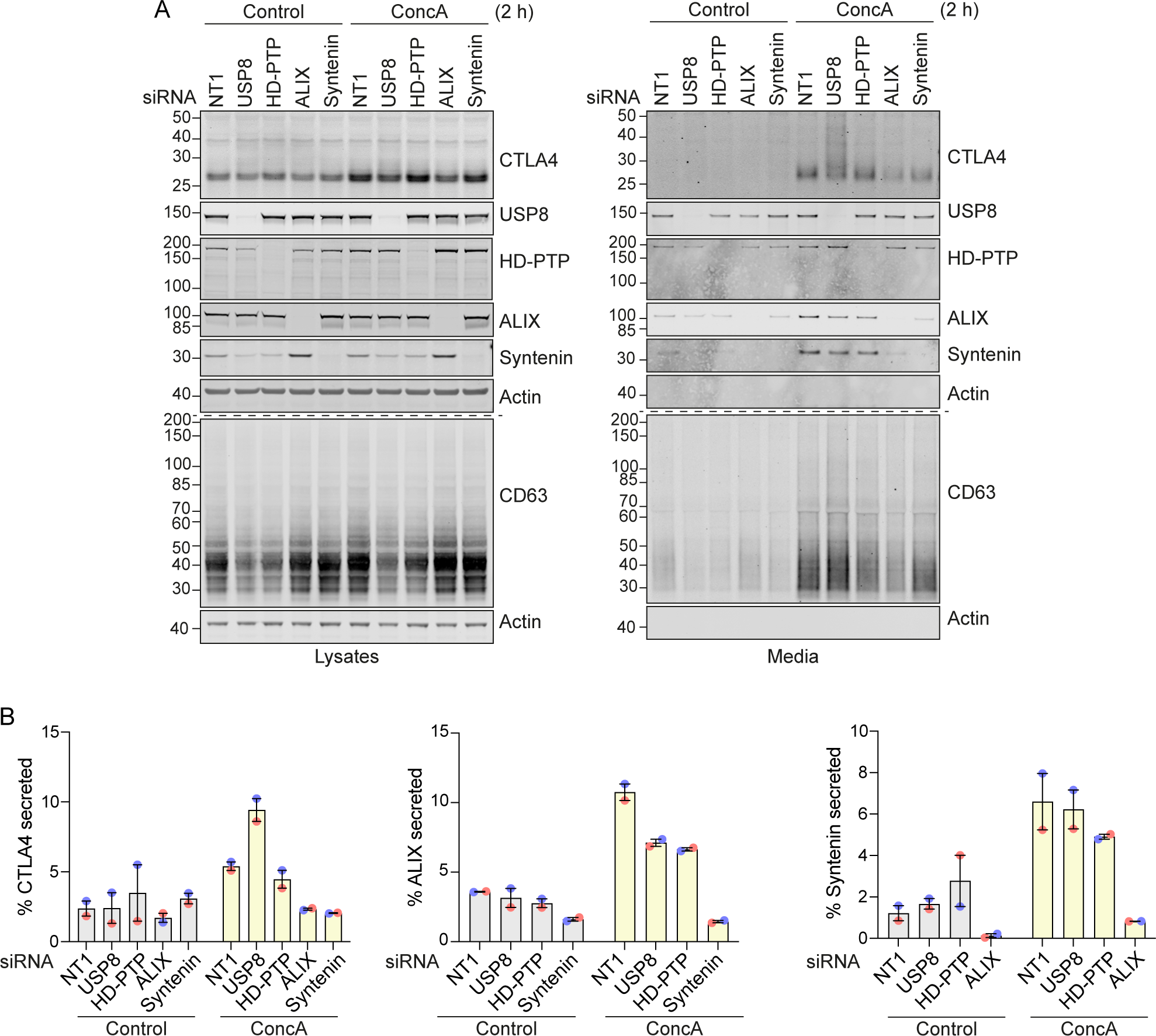
Ubiquitylated CTLA4 is secreted via exosomes from USP8-depleted cells. **A.** A2058 cells were transfected for 72 h with non-targeting (NT1) or USP8, HD-PTP, ALIX and Syntenin siRNAs. Lysates were collected and the cultured supernatant subjected to TCA precipitation before analysis by SDS-PAGE and western blot. **B.** Quantification of CTLA4 secreted into the media for data represented in (A). Individual data points from two independent, colour-coded experiments are shown. Error bars indicate the range.

**Figure S4.**
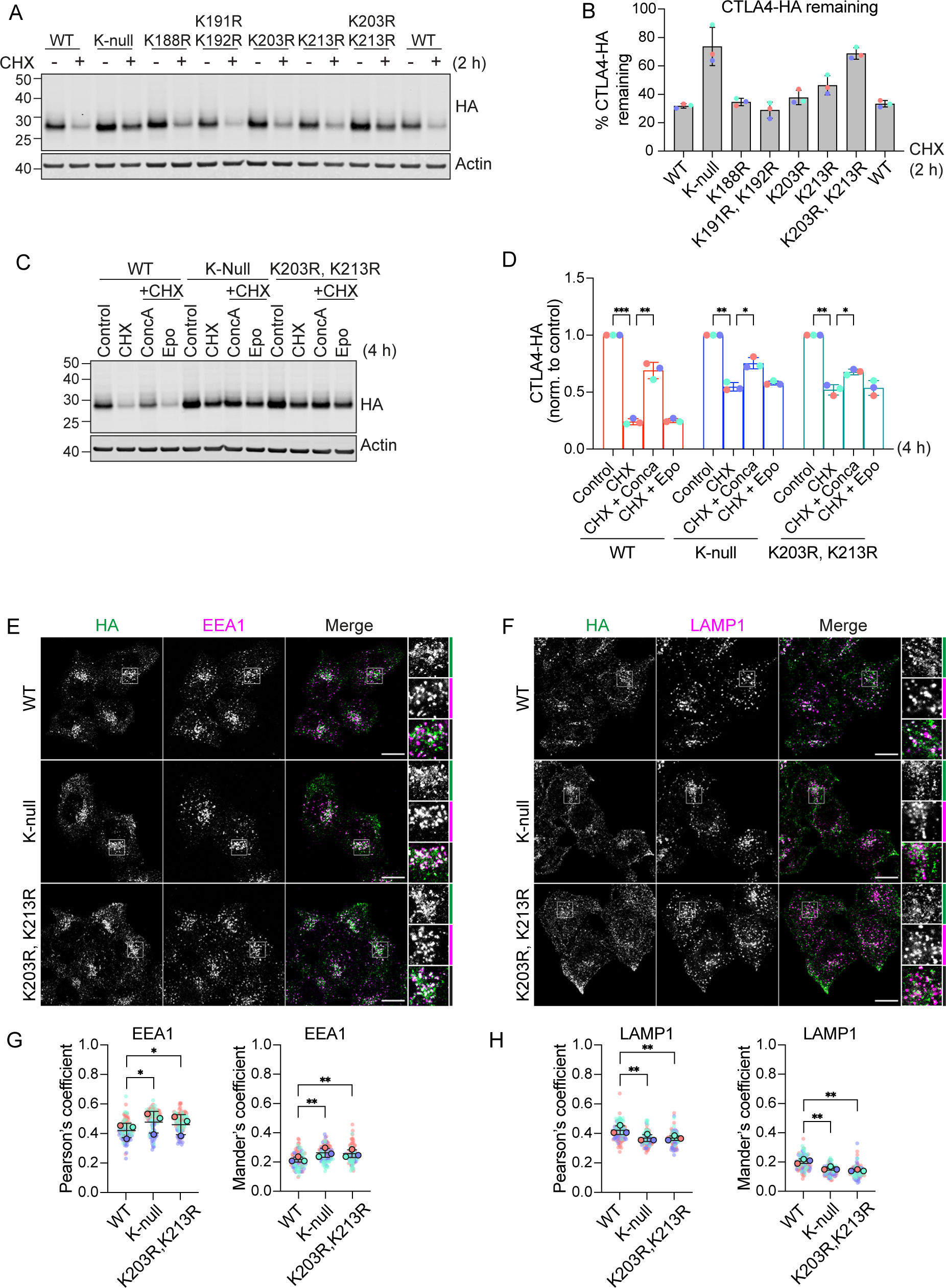
Mutation of K203 and K213 delays CTLA4 turnover. **A.** Representative western blots showing increased stability of CTLA4-HA K-null and K203R,K213R mutants. HeLa S3 Flp-In CTLA4-HA WT and indicated lysine mutants were treated with Cycloheximide (CHX) for 2 h before lysis. **B.** Quantification of CTL4-HA remaining following CHX treatment. Individual data points from three independent, colour-coded experiments are shown. Error bars show standard deviation. **C.** Representative western blots of HeLa S3 Flp-In CTLA4-HA WT, K-null and K203R, K213R double lysine mutants treated for 4 h with Cycloheximide (CHX) alone or together with Concanamycin A (ConcA) or Epoxomicin (Epo) prior to lysis. **D.** Quantification of CTLA4-HA remaining normalised to control treated cells for data represented in (C). Individual data points from three independent, colour-coded experiments are shown. Error bars show standard deviation. Two-way ANOVA multiple comparisons with uncorrected Fisher’s LSD. *p<0.05, **p<0.01, ***p<0.001. **E,F.** Representative confocal images of CTLA4-HA WT, K-null and K203R,K213R mutants co-stained with EEA1 (E) or LAMP1 (F). Scale bar = 15 µm. **G,H.** Co-localisation analysis of CTLA4-HA WT, K-null and with EEA1 (G) and LAMP1 (H) . Graphs show Pearson’s coefficients or Mander’s coefficient. Error bars indicate standard deviation for three independent, colour-coded experiments. Opaque circles with dark outline correspond to the mean value from each experiment. One-way ANOVA with Dunnett’s multiple comparisons test. *p<0.05, **p<0.01.

**Figure S5.**
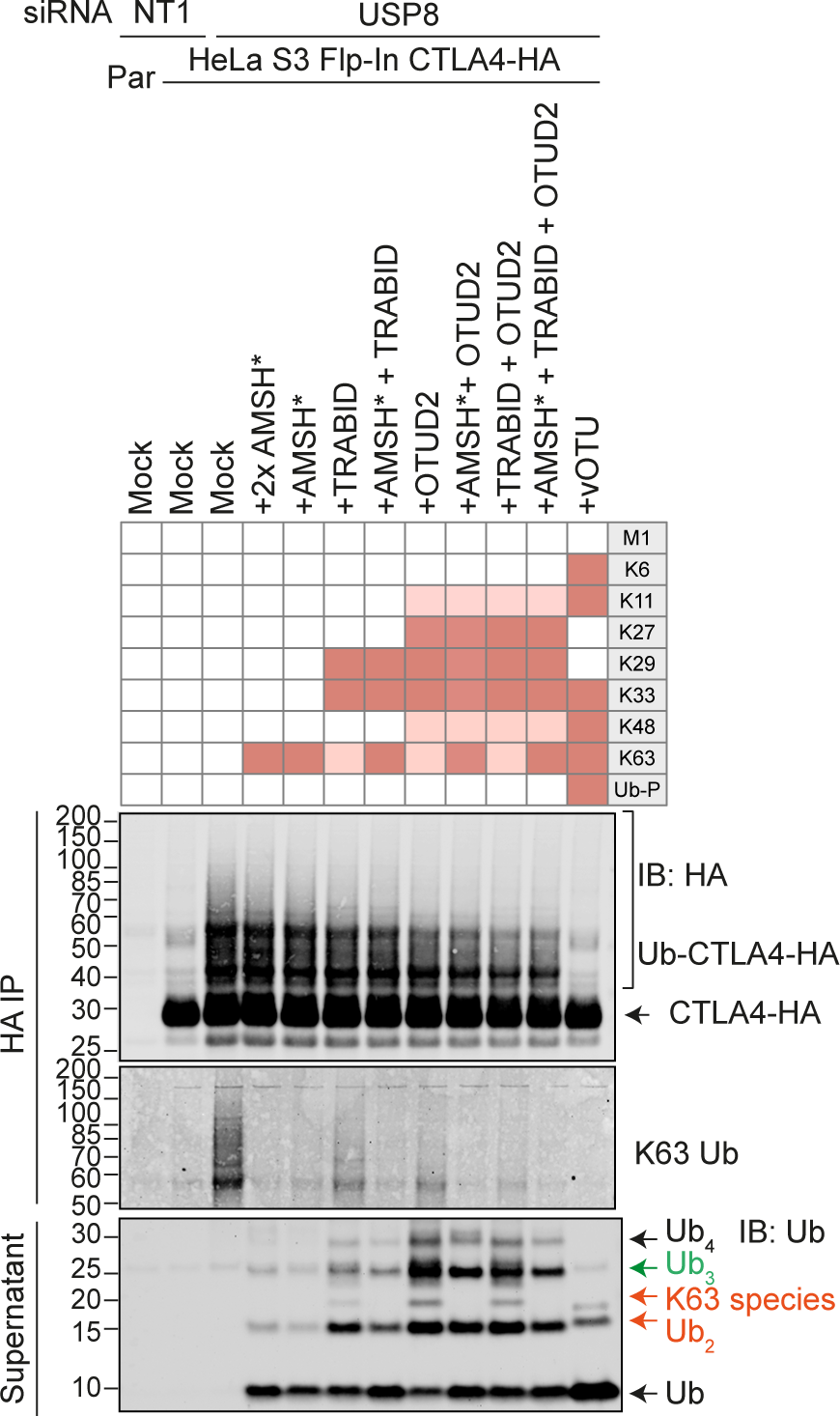
Combinatorial UbiCRest analysis of CTLA4-HA. Representative western blot of UbiCRest analysis of CTLA4 using a combination of linkage-specific DUBs. HeLa S3 Flp-In CTLA4-HA cells were transfected for 72 h with NT1 or USP8 siRNA prior to lysis. CTLA4-HA was immunoprecipitated using anti-HA magnetic beads and incubated for 1 h at 37°C with the indicated DUBs. Supernatants containing ubiquitin species released by DUBs were collected and analysed in parallel with CTLA4-HA eluted from the beads. Specific activities of DUBs as reported in the literature are depicted in the key above the blot. Ub-P indicates ability to cleave proximal Ubiquitin.

